# Multisensory inputs control the regulation of time investment for mating by sexual experience in male *Drosophila melanogaster*

**DOI:** 10.1101/2022.09.23.509131

**Authors:** Seung Gee Lee, Changku Kang, Baraa Saad, Khoi-Nguyen Ha Nguyen, Adrian Guerra-Phalen, Dorothy Bui, Al-Hassan Abbas, Brian Trinh, Ashvent Malik, Mahdi Zeghal, Anne-Christine Auge, Md Ehteshamul Islam, Kyle Wong, Tiffany Stern, Elizabeth Lebedev, Dongyu Sun, Hongyu Miao, Zekun Wu, Thomas N. Sherratt, Woo Jae Kim

## Abstract

Males have finite resources to spend on reproduction. Thus, males rely on a ‘time investment strategy’ to maximize their reproductive success. For example, male *Drosophila melanogaster* extends their mating duration when surrounded by conditions enriched with rivals. Here we report a novel form of behavioral plasticity whereby male fruit flies exhibit a shortened duration of mating when they are sexually experienced; we refer to this plasticity as ‘shorter-mating-duration (SMD)’. SMD is a plastic behavior and requires sexually dimorphic taste neurons. We identified several neurons in the male foreleg and midleg that express specific sugar, pheromone and mechanosensory receptors. Using a cost-benefit model and behavioral experiments, we further show that SMD behavior exhibits adaptive behavioral plasticity in male flies. Thus, our study delineates the molecular and cellular basis of the sensory inputs required for SMD; this represents a plastic interval timing behavior that could serve as a model system to study how multisensory inputs converge to modify interval timing behavior for improved adaptation.

**ONE SENTENCE SUMMARY:** Male flies use information derived from their previous sexual experiences from multiple sensory inputs to optimize their investment in mating.

## INTRODUCTION

From basic behaviors to complicated decisions, all animals have to make choices throughout their life to maximize their utility function [1]. The reproductive success of a male animal depends predominantly on how many of its sperm are successful in fertilizing eggs [2]. Males have a finite resource to spend on reproduction [3] and must make choices throughout their life to optimize how their resources are utilized [4]. For example, males that invest a long period of time for mating might expose themselves to the action of predators or various environmental hazards, thereby losing their competitiveness. In this regard, the ‘time investment strategy’ (the optimum allocation of time spent on given activities to achieve maximal reproductive success)’ is crucial for males. Female guarding has commonly evolved as a time investment strategy for males [5].

Recent studies have revealed that male *D. melanogaster* shows wide variation in terms of their level of interest in females, thus providing evidence that males have also evolved to mate selectively [6]. When mating opportunities are constrained, males that show a preference for more fecund females will benefit directly by increasing the number of offspring they produce [7]. The selective mating investment exhibited by male *D. melanogaster* may have evolved for several reasons. First, sexual activity reduces the lifespan of males (Partridge and Farquhar, 1981) due to costs arising from vigorous courtship (Cordts and Partridge, 1996), the production of ejaculates [8] and possibly also due to immunosuppression [9]. Second, repeated mating by males within a 24 h period depletes limiting components of the ejaculate [10]. Third, the quality of potential female mates is highly variable [11].

Behavioral plasticity is advantageous when specific aspects of the environment (e.g., the intensity of socio-sexual encounters) are prone to rapid and unpredictable variation [12]. The best-studied example of plastic behavioral responses in males is ‘longer-mating-duration (LMD)’ in which exposure to rivals before mating increases investment through mating duration [13, 14].

Here, we report a novel form of plastic behavior in male *D. melanogaster* with regards to time investment in mating. We found that sexually experienced *Drosophila* males exhibit this plastic behavior by limiting their investment in copulation time; we refer to this mechanism as ‘shorter-mating-duration (SMD)’.

## RESULTS

To investigate how sexual experience affects the mating duration of male *D. melanogaster*, we introduced virgin females to group-reared males one day before the assay (this condition is referred to as ‘experienced’ hereafter) and compared their mating duration with group-reared males that had never encountered sexual experience (this condition is referred to as ‘naïve’ hereafter) (Fig. 1A). We found that the mating duration of various wild-type and *w^1118^* naïve males are significantly longer than that of sexually experienced males (Fig. 1B-D, Fig. S1A), thus suggesting that the effect of the *white* mutant genetic background was not evident unlike that of LMD behavior as reported previously [14]. *Drosophila simulans*, the sibling species of *D. melanogaster* also exhibits SMD, thus suggesting that SMD is conserved between close species of *D. melanogaster* (Fig. S1B).

**Fig. 1.**
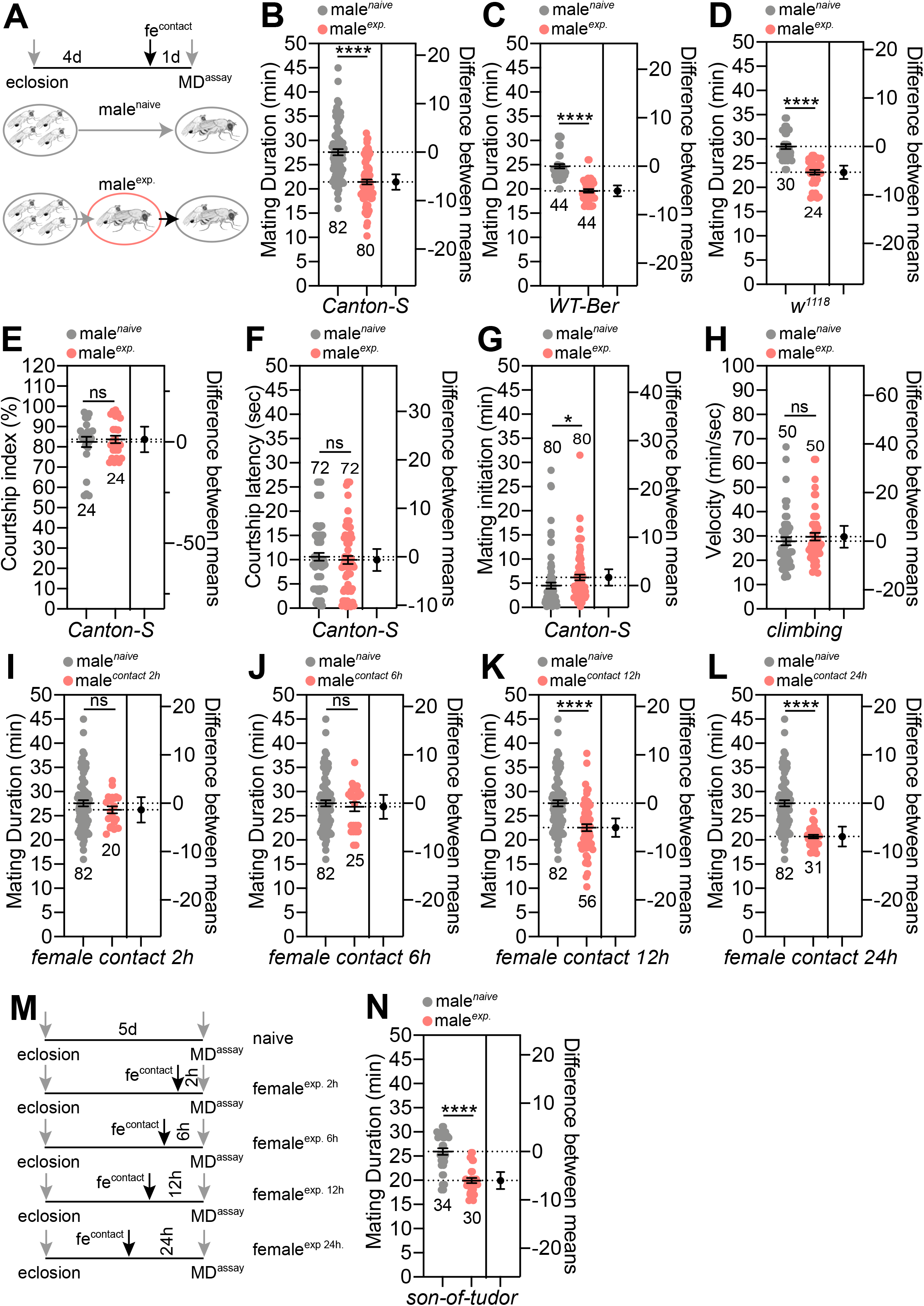
General characteristics of ‘shorter-mating-duration (SMD)’ behavior. (A) Naïve males were kept for 5 days after eclosion in groups of 4 males. Experienced males were kept for 4 days after eclosion in groups then experienced with 5 virgin females 1 day before assay; for detailed methods, see the **EXPERIMENTAL PROCEDURES**. (B) Mating duration (MD) assays of Canton-S (CS), (C) WT-Berlin, and (D) *w^1118^* males. Light grey dots represent naïve males and pink dots represent experienced ones. (E) Courtship index of naïve and experienced males. See the **EXPERIMENTAL PROCEDURES** section for detailed methods. (F) Courtship latency of naïve and experienced males. See the **EXPERIMENTAL PROCEDURES** section for detailed methods. (G) Mating initiation time of naïve and experienced males. (H) The locomotion of naïve and experienced male flies were quantified as velocity by a climbing assay paradigm. (I-L) MD assays of CS males with different exposure time with females. Each group of males was reared with females for (I) 2 h, (J) 6 h, (K) 12 h or (L) 24 h. (M) A diagram showing the results of MD assays of CS males with different exposure times with females. (N) MD assays for *son-of-tudor* mutants. Genotypes are described as in a previous report [15]. Dot plots represent the MD of each male fly. The mean value and standard error are labeled within the dot plot (black lines). Asterisks represent significant differences, as revealed by the Student’s *t* test (* *p<0.05*, ** *p<0.01*, *** *p<0.001*). The same notations for statistical significance are used in other figures. Number signs represent significant differences, as revealed by Dunn’s Multiple Comparison Test (^#^ *p<0.05*). The same symbols for statistical significance are used in all other figures. See the **EXPERIMENTAL PROCEDURES** for a detailed description of the statistical analysis used in this study.

To test whether fatigue causes SMD behavior, we examined other behavioral repertoires of naïve and experienced male flies, such as courtship index, courtship latency, copulation latency and locomotion; there was no significant difference between experienced and naïve males (Fig. 1E-H, Fig.S1C-D). Thus, we conclude that potential fatigue from repetitive sexual experiences is not a causative factor for SMD behavior.

To determine the time required by males to be exposed to females in order to induce SMD behavior, we varied the exposure time of males to females and found that males significantly reduced their mating duration when their exposure to females lasted for longer than 12 h but not for less than 6 h, thus suggesting that SMD requires chronic exposure to females for longer than 6 h (Fig. 1I-M). To determine whether SMD is a reversible behavior, we separated males from females after 24 h or 48 h of sexual experience and then tested these males in a mating duration assay. We found that separating experienced males from females for 24 h was sufficient to restore the MD to the level of naïve males (Fig. S1E-H), thus suggesting that SMD is plastic and dependent on sexual experience with females but can change over time.

To confirm the lack of effect of sperm depletion on SMD behavior, we depleted sperm prior to MD assays and found that sperm depletion did not affect SMD behavior (Fig. 1I-M). We also tested the *son-of-tudor* males that lack germ cells and are therefore devoid of sperm [15]; we found that the *son-of-tudor* males also exhibited SMD (Fig. 1N). Consistent with a previous report [16], these data suggest that sperm depletion does not cause SMD behavior in male *D. melanogaster*.

Next, to identify the sensory modalities that modulate SMD behavior, we tested multiple mutants with defects in various sensory modalities [14, 17]. By using constant dark conditions (Fig. 2A) and several mutants with impaired vision (*GMR-Hid* in Fig. 2B; *ninaE^17^* in Fig. 2C), impaired olfaction (*Orco^1^/Orco^2^* in Fig. 2D), impaired gustation (*GustD^x6^* in Fig. 2E) and impaired auditory ability and mechanosensation (*iav^1^* in Fig. 2F), we concluded that gustatory, auditory and mechanosensory pathways are involved in generating SMD behavior but not visual or olfactory pathways.

**Fig. 2.**
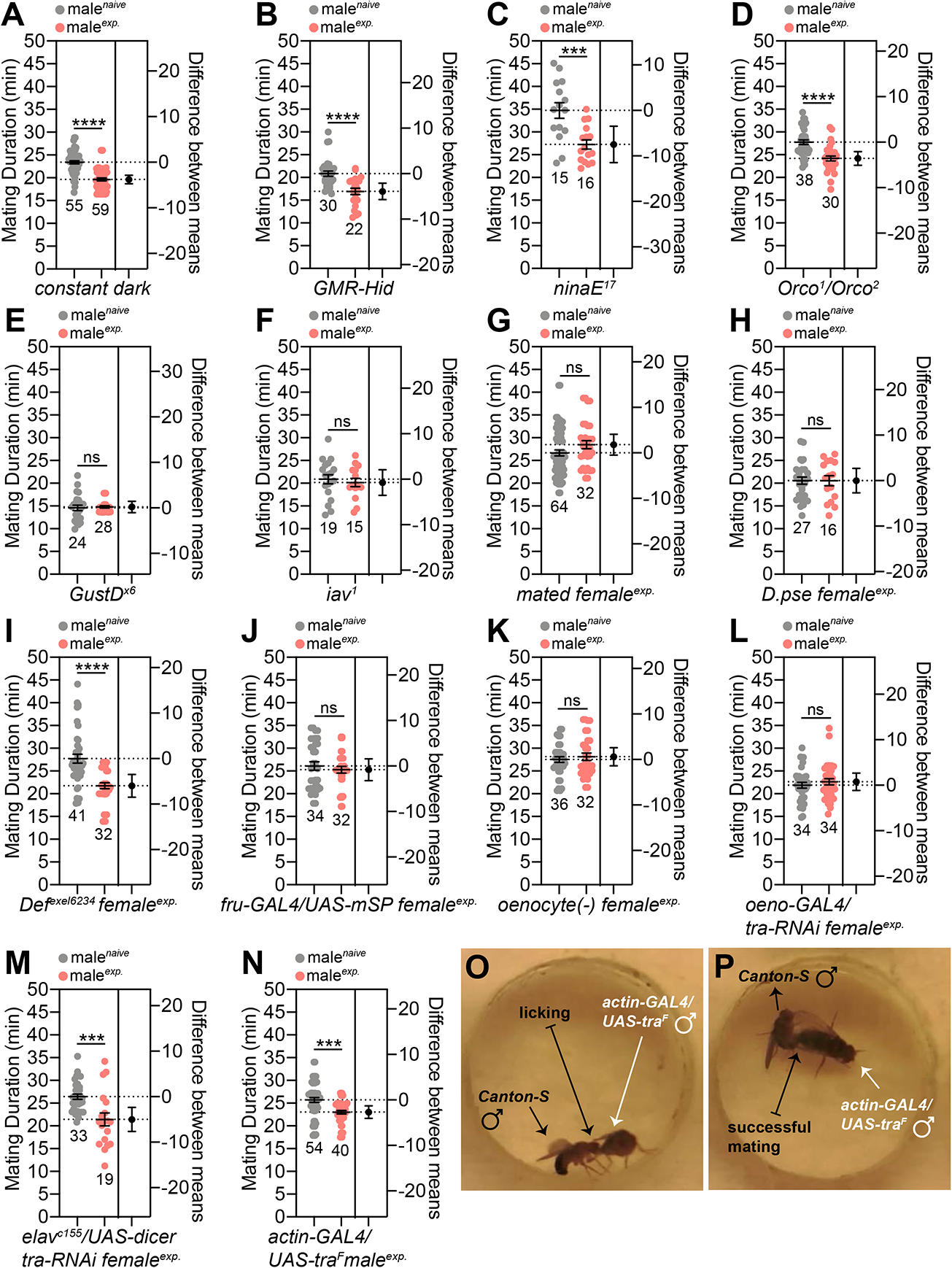
Sensory inputs required for inducing SMD behavior. (A) To test whether the vision is required for SMD, CS males were reared and sexually experienced in constant dark for 5 days (dark). (B) MD assays of *GMR-Hid* males, and blind animal. (C) MD assays of *ninaE^17^* mutant males animal lacking the opsin R1-6 photoreceptors [81]. (D) MD assay of *Orco^1^*/*Orco^2^* trans-heterozygote mutant males with defects in olfaction [82]. (E) MD assays of *GustD^x6^* mutant males showing aberrant responses to sugar and NaCl [83]. (F) MD assays of *iav^1^* males, the auditory mutant [84]. (G) MD assays of CS males exposed to sexually experienced females 1 day before assay. To generate mated females, 4-day-old 10 CS virgin females were placed with 5-day-old 20 CS males for 6 hours and then transferred to an empty vial. These females were used for experienced females 1 day after separation. (H) MD assay of CS males experienced with *D. pseudoobscura* females. (I) MD assay of CS males experienced with *Df^exel6234^* females, a deficiency strain that lacks the expression of the sex-peptide receptor (SPR) [85]. (J) MD assays of CS males experienced with virgin females behaving as mated females. To make virgin females behave as mated females, flies expressing *UAS-mSP* (a membrane bound form of male sex-peptide) were crossed with flies expressing *fru-GAL4* driver, as described previously [18, 86]. (K) MD assays of CS males experienced with oenocyte-deleted females. To generate oenocyte-deleted females, virgin flies expressing *UAS-Hid/* crossed with flies expressing *tub-GAL80ts, oeno-GAL4* males; then the female progeny were kept in 22℃ for 3 days. Flies were moved to 29℃ for 2 days before assay to express *UAS-Hid/rpr* and kill the oenocytes in these females. The *oeno-GAL4* (*PromE(800)-GAL4*) was described previously [87]. (L) MD assays of CS males exposed to oenocyte-masculinized females. To generate oenocytes-masculinized females, flies expressing *UAS-tra-RNAi* were crossed with *oeno-GAL4* driver. (M) MD assays of CS males exposed to pan-neuronally masculinized females. To generate pan-neuronally masculinized females, flies expressing *UAS-tra-RNAi* were crossed with *elav^c155^* driver. (N) MD assays of CS males exposed to feminized females. To generate feminized males, flies expressing *actin-GAL4* were crossed with flies expressing *UAS-tra^F^*. (O) CS male courting with a feminized male and showing licking behavior, leading to successful mating (P).

Next, we attempted to identify the physiological cues from females that play important roles in the induction of SMD behavior in males. To do this, we used various genotypes of females as experienced sexual partners. Mated females and *Drosophila pseudoobscura* females did not induce SMD, thus suggesting that cues originate from virgin *D. melanogaster* females (Fig. 2G-H). Excessive mating with sex-peptide receptor (SPR) mutant females during the experimental period, which exhibit a delayed post-mating response [18], had no additional effect on SMD (Fig. 2I). Virgin females behave like mated females by expressing a membrane-bound form of male sex-peptide in *fruitless*-positive neurons [18]. Males that were experienced with these females did not show SMD, thus suggesting that both cues from females and successful copulation are required for SMD (Fig. 2J).

We produced odorless and tasteless females by killing female oenocytes *(oenocyte(-)*) and females that produced a male odor *via* the masculinization of female oenocytes (*oeno-GAL4/tra-RNAi*). Males that had experience with these females did not show SMD, thus suggesting that female-specific pheromones produced by oenocytes are important cues for SMD (Fig. 2K-L). However, males experienced with females which contained masculinized neurons showed intact SMD, thus suggesting that female forms of odor, and not female forms of neural circuits, are critical for inducing SMD behavior (Fig. 2M). Interestingly, feminized males, created by overexpressing the female form of the *tra2* protein driven by a broad *GAL4* driver, can provide the cues required for SMD, thus suggesting that developmental phenotypes that are regulated by *tra2* can provide both cues from females and successful copulation that are sufficient to induce SMD (Fig. 2N). By tracking videos of the mating assay, we were able to confirm that males exhibited a full repertoire of courtship behavior and mated successfully with oenocyte-masculinized females (Fig. S2A-F) and feminized males (Fig. 2O-P, Fig. S2G-H), thus suggesting that these experienced partners can provide a mating drive for male *D. melanogaster*. We also found that SMD was completely normal even when an oenocyte-masculinized female (Fig. S2I) was used for assay partners, thus suggesting that SMD is independent of the genotypes of the assay partners used for mating duration assays. Collectively, these data suggest that both sexual experience and female *D. melanogaster*-specific odor (produced in the oenocytes) are required to induce SMD behavior.

In flies, taste and touch signals are primarily conveyed to the brain by sensory neurons in the legs and mouthparts. To understand how sensory information for SMD is mediated *via* the legs or proboscis, we first tested the SMD behavior of males for which each pair of legs had been removed; we found that the foreleg is critical for generating SMD behavior (Fig. 3A-C). When we carefully watched the position of each pair of legs during mating, we found that the male’s foreleg touches the female body most of the time during mating; the midleg only partially touches the female body while the hind leg does not touch the female at all (Fig. 3D-G). The point at which the male’s leg touched the female body was mostly the tarsus, an area that is known to recognize taste [19] and pheromones [20] *via* chemoreception (Fig. S3A). Although we cannot rule out the role of the proboscis, wings and other unidentified taste organs in the reception of stimuli for SMD behavior, our present results suggest that the male’s foreleg is the major sensor for SMD behavior.

**Fig. 3.**
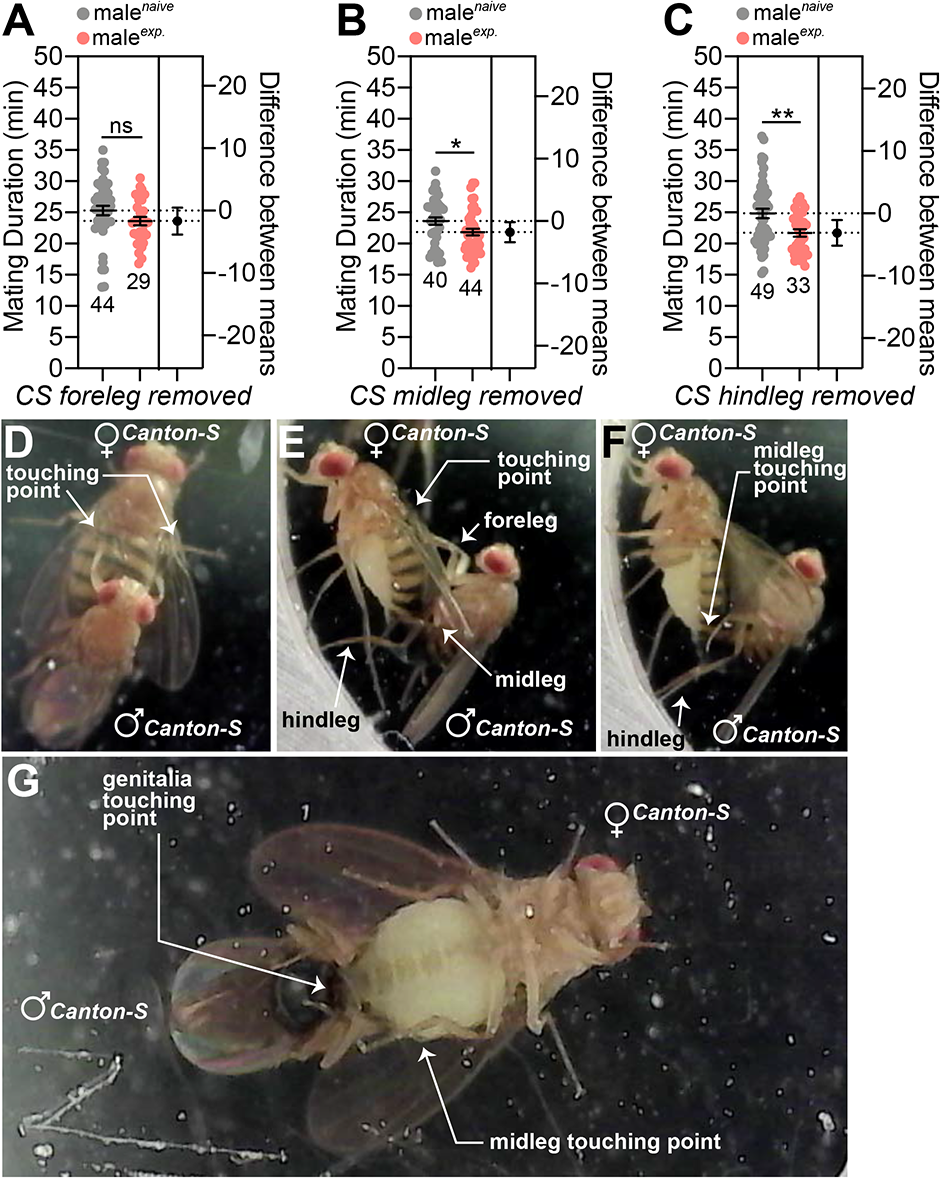
The male foreleg is crucial to detect the sensory inputs to induce SMD behavior. (A) MD assays of CS males in which the foreleg, (B) midleg, or (C) hindleg were removed 1 day before assay. Forelegs, midlegs, or hindlegs of 4-day-old males were removed by surgery and then treated as naïve or experienced for 1 d. (D) Dorsal view of the mating posture of CS males and females. The touching point of the male foreleg is marked with a white arrow. (E) Lateral view of the mating posture of CS males and females. The touching points of the male foreleg and midleg are marked with a white arrow. (F) Lateral view of the mating posture of CS males and females. The touching point of the male midleg is marked with a white arrow. (G) Ventral view of the mating posture of CS males and females. The touching points of the male midleg and genitalia are marked with a white arrow.

Of the various gustatory receptors, *Gr5a* marks cells that recognize sugars and mediate taste acceptance, whereas *Gr66* a is a salt receptor in Drosophila that recognizes bitter compounds and mediates avoidance. *Gr5a* and *Gr66a* are expressed in different cells in a sensillum of the foreleg and exhibit different sensory projections into the central brain region (Fig. 4A-B). We found that male flies with ablated *Gr5a*-positive neurons that mediate sweet-taste detection did not exhibit SMD behavior while male flies lacking *Gr66a*-positive neurons that mediate bitter-taste detection exhibited normal SMD (Fig. 4C-D). SMD was also impaired when we inhibited synaptic transmission *via* the expression of *TNT* in *Gr5a*-positive neurons but not in *Gr66a*-positive ones in an adult-specific manner by using *tub-GAL80^ts^* (Fig. 4E-F). The inactivation or hyperexcitation of *Gr5a*-positive neurons, but not *Gr66a*-positive neurons, by expressing the *KCNJ2* potassium channel or *NachBac* bacterial sodium channel in an adult-specific manner using *tub-GAL80^ts^*, also resulted in impaired SMD (Fig. 4G-J). These data and genetic background control data (Fig. S4A-D) suggest the cell populations of gustatory cells that mediate acceptance signals are associated with SMD behavior and that these *Gr5a*-positive neuronal populations and their neuronal activities are required for SMD.

**Fig. 4.**
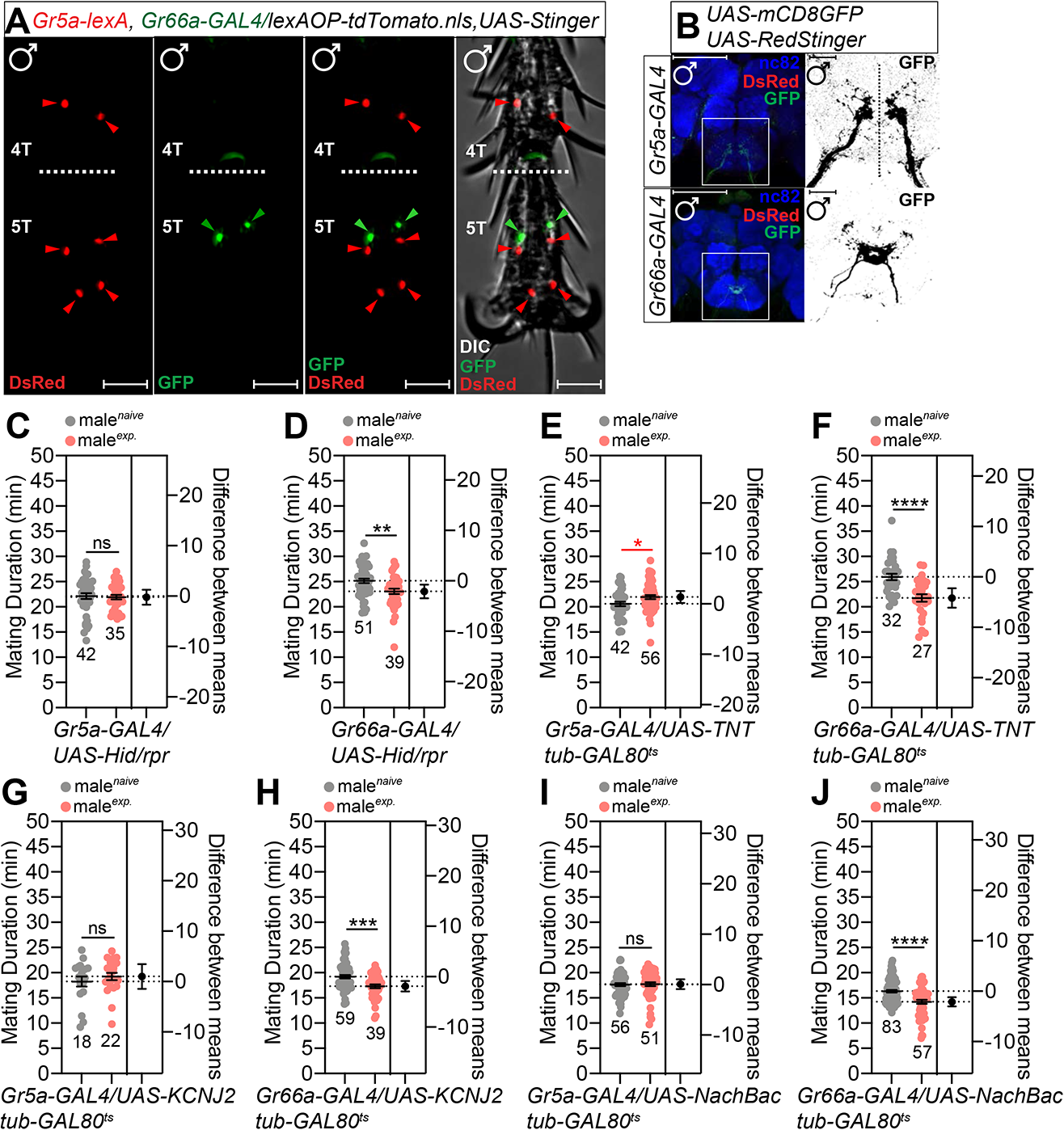
Gr5a-positive sugar cells are important for inducing SMD behavior in the male foreleg. (A) 4T and 5T of the male foreleg of flies expressing *Gr5a-lexA* and *Gr66a-GAL4* drivers together with *lexAOP-tdTomato* and *UAS-Stinger* were imaged live under a fluorescent microscope. Red arrows indicate *Gr5a-* positive neurons and green arrows indicate *Gr66a*-positive neurons. Scale bars represent 50 μm. (B) Brains of flies expressing *Gr5a-GAL4 or Gr66a-GAL4* together with *UAS-mCD8GFP, UAS-RedStinger* were immunostained with anti-GFP (green), anti-DsRed (red) and nc82 (blue) antibodies. Scale bars represent 100 μm. The right panels indicate magnified regions of the left panels that are presented as a grey scale to clearly show the axon projection patterns of gustatory neurons in the adult sub-esophageal ganglion (SOG) labeled by *GAL4* drivers. (C-D) MD assays for *GAL4* driven cell death which labelled (C) sweet cells (*Gr5a*) or (D) bitter cells (*Gr66a*) using *UAS-Hid/rpr*. (E-F) MD assays of (E) *Gr5a-* or (F) *Gr66a-GAL4* drivers for the inactivation of synaptic transmission *via* the expression of *UAS-TNT* transgene together with the *tub-GAL80^ts^*, such that *UAS-TNT* expression could be triggered by temperature shifts were crossed with flies expressing *tub-GAL80^ts^* (G-J) electrical silencing or hyperexcitation of *Gr5a*-positive neurons abolished SMD behavior. Flies expressing (G-H) potassium channel *UAS-KCNJ2* or (I-J) bacterial voltage-gated sodium channel *UAS-NachBac* together with the *tub-GAL80^ts^*, such that *UAS-KCNJ2* or *UAS-NachBac* expression could be triggered by temperature shifts, were crossed with flies expressing (G and I) *Gr5a-* or (H and J) *Gr66a-GAL4* drivers. Flies were reared at 29℃ for the first 2 days to strongly induce *UAS-KCNJ2* or *UAS-NachBac* expression and then transferred to 25℃ for the last 3 days for the mild induction of *UAS-KCNJ2* or *UAS-NachBac* transgenes.

In addition, we found that *Gr5a*-positive cells were abundantly localized in the tarsus from tarsomeres 2 (2T) to tarsomeres 5 (5T) (Fig. S4E). We also found that males have more Gr5a-positive cells than females (Fig. S4F). On average, males had 10 ± 1 neurons in the tarsus (4 cells in 5T, 2 ± 1 cells in 4T, 1 ± 1 cells in 3T, 2 cells in 2T and no cells in 1T) and 0 ± 1 cells in the tibia; however, females had 6 cells in the tarsus (4 cells in 5T and 2 cells in 4T) (Fig. S4E-F). These data suggest that *Gr5a*-positive cells show sexual dimorphism and might have a male-specific function to generate SMD.

The sexual dimorphism of sensory structure and function generates neural circuitries that are important for gender-specific behaviors. In *Drosophila*, *fruitless (fru)* is an essential neural sex determinant that is responsible for male-specific behavior [21]. To determine whether sexually dimorphic sensory neurons are involved in SMD, we used intersectional methods to genetically dissect approximately 1500 *fru* neurons into smaller subsets. We used a combination of the *fru^FLP^* allele that drives FLP-mediated recombination specifically in *fru* neurons with *UAS>stop>X* genotype (X represents various reporters or effector transgenes) to express a *UAS* transgene in only those cells that were labeled by the *GAL4* driver and were also *fru*-positive; this was controlled by the FLP-mediated excision of the stop cassette (*>stop>*).

We found that the sensory projections of a subset of *Gr5a*-positive neurons, but not *Gr66a*-positive neurons, were positive for *fruitless,* an essential neural sex-determinant that is responsible for male-specific behaviors [21] (Fig. 5A and Fig. S5A). To test whether the small subset of *fru*-positive Gr5a cells is involved in SMD, we expressed tetanus toxin light chain (*UAS>stop>TNT_active_)* with *Gr5a-* or *Gr66a-GAL4* drivers along with *fru^FLP^* to inhibit synaptic transmission in sexually dimorphic subsets of *fru*-positive cells. We found that SMD was abolished when *UAS-TNT* was expressed only in male-specific *Gr5a*-positive neurons (Fig. 5B-C). As a control, we found that SMD was unaffected when we used each of these *GAL4* drivers in combination with *UAS>stop>TNT_inactive_* to express an inactive form of the tetanus toxin light chain (Fig. 5D-E). The systemic expression of a female form of *tra* cDNA (*UAS-tra^F^*) in a male during development is known to lead to the expression of female characteristics [22]. We found that SMD was eliminated by the feminization of *Gr5a-GAL4* labeled cells but not by the expression of *UAS-tra^F^* in Gr66a-positive neuronal subsets (Fig. 5F-G), thus suggesting that the feminization of *Gr5a*-positive neurons nullifies the male-specific sensory function of those cells to detect sensory inputs for SMD behavior. Together with genetic background control experiments (Fig. S5B-D), these data suggest that SMD requires the male-specific role of a subset of *Gr5a*-positive neurons.

**Fig. 5.**
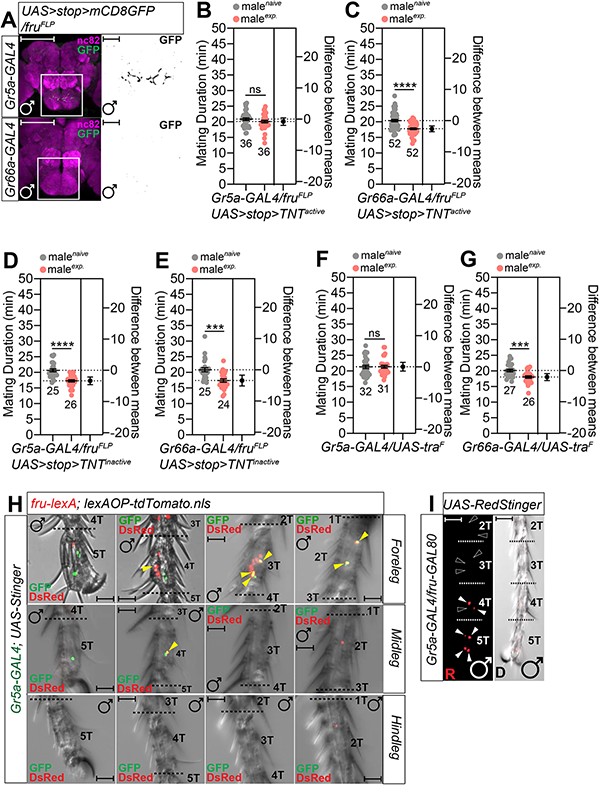
FRU-positive subsets of sugar sensing neurons in the leg are required to generate SMD. (A) Brains of male flies expressing *Gr5a-GAL4* or *Gr66a-GAL4* together with *UAS>stop>mCD8GFP; fru^FLP^* were immunostained with anti-GFP (green) and nc82 (magenta) antibodies. Scale bars represent 100 μm in the colored panels and 10 μm in the grey panels. White boxes indicate the magnified regions of interest presented in the right panels. The right panels are presented as a grey scale to clearly show the axon projection patterns of gustatory neurons in the adult sub-esophageal ganglion (SOG) labeled by *GAL4* drivers. (C-D) MD assays of (C) *Gr5a-* and (D) *Gr66a-GAL4* drivers for the inactivation of synaptic transmission of *fru*-specific neurons among each *GAL4*-labelled neuron via *UAS>stop>TNT_active_; fru^FLP^*. (D-E) Control experiments of (B-C) with the inactive form of *UAS-TNT* using *UAS>stop>TNT_inactive_; fru^FLP^*. (F-G) MD assays for (F) *Gr5a-* and (G) *Gr66a-GAL4* drivers for the feminization of neurons *via UAS-tra^F^*. (H) Male foreleg, midleg and hindleg tarsus of flies expressing *fru-lexA* and *Gr5a-GAL4* drivers together with *lexAOP-tdTomato* and *UAS-Stinger* were imaged live under a fluorescent microscope. Red arrows indicate *Gr5a-*positive neurons and green arrows indicate *Gr66a*-positive neurons. Scale bars represent 50 μm. (I) Male foreleg of flies expressing *Gr5a-Gal4* together with *fru-GAL80*. White arrows indicate *Gr5a*-positive and *fru*-negative neurons. Dotted white arrows indicate missing neurons by adding *fru-GAL80*, as shown in Fig. S4F.

By using the genetic intersectional method [23], we found that the male foreleg contains 5 - 6 *Gr5a*- and *fru*-positive cells in the tarsus (1 in 4T, 2 - 3 in 3T and 2 in 2T) while the midleg contains 1 (1 in 4T) (Fig. 5H). However, we could one of these cells in the male proboscis (Fig. S5E). We also confirmed the number and position of *Gr5a*-expressing *fru*-positive cells using *fru-GAL80* combined with *Gr5a-GAL4*, as shown in Fig. 5H (Fig. 5I). Together with the data arising from leg removal experiments (Fig. 3), these data suggest that *Gr5a*-expressing male-specific sensory cells in the male leg provide the major sensory input for SMD generation.

Next, we asked whether sugar receptors in the sexually dimorphic sugar sensory neurons are involved in the generation of the sensory input pathways that generate SMD. Sugars are the main group of chemicals underlying sweet taste and provide essential nutritional value for many mammals and insects [24]. Sweet taste in *D. melanogaster* is mediated by eight, closely related gustatory genes: *Gr5a*, *Gr61a*, and *Gr64a*-*Gr64f* [25]. The *Gr5a^lexA^* allele refers to the Gr5a gene replaced by the mini-white transgene [25] results in a lack of SMD, thus suggesting that *Gr5a* itself is an important receptor for generating SMD (Fig. 6A). We knocked down all known sugar receptors in *fru*-positive cells using a *fru-GAL4* driver and found that only *Gr5a* and *Gr64f* are important for the generation of SMD in male-specific *fru*-positive cells (Fig. 6B-D, Fig. S6A-G).

**Fig. 6.**
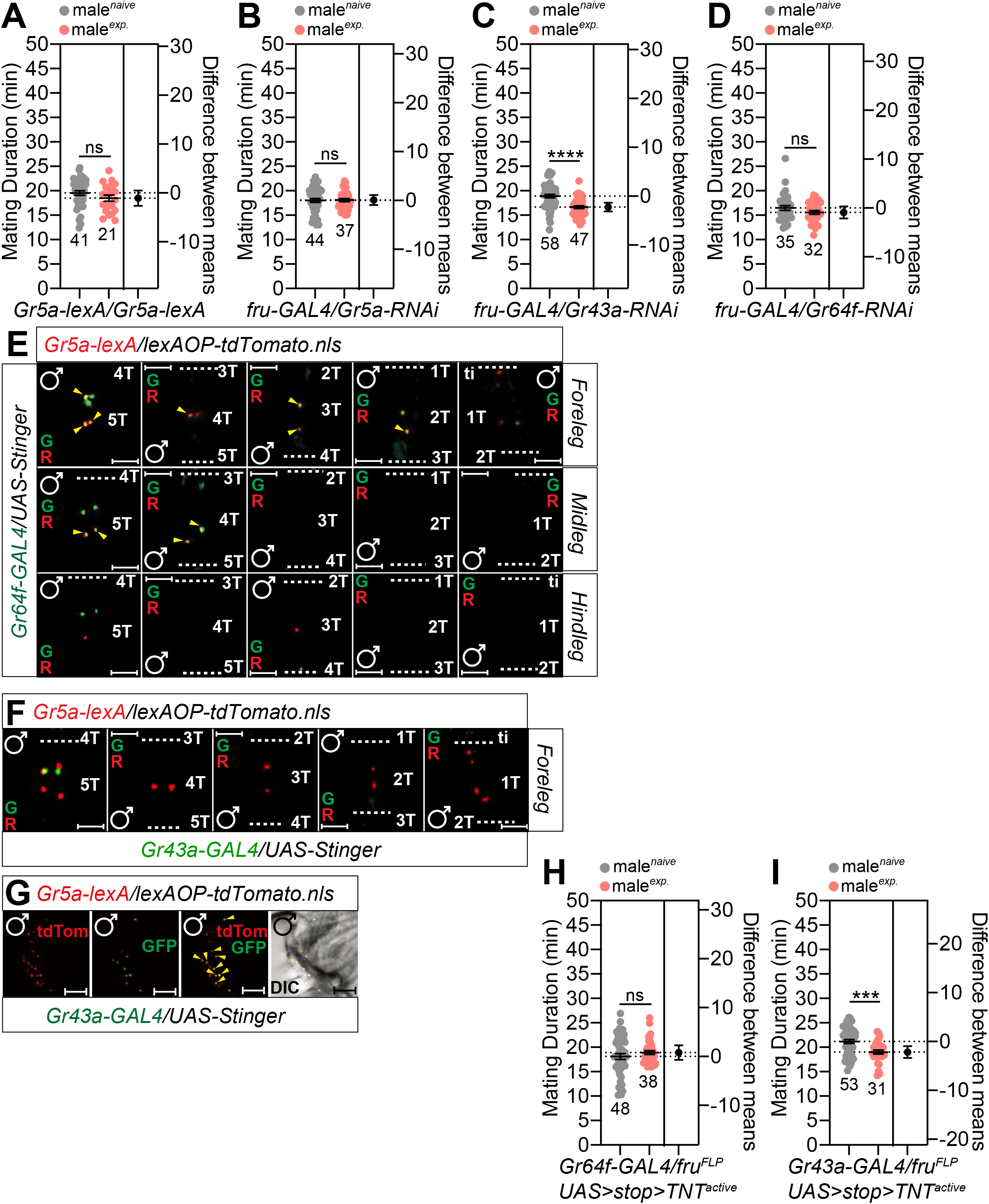
Specific sugar receptors in the male foreleg are critical for inducing SMD behavior. (A) MD assay of *Gr5a-lexA* homozygote males in which the *Gr5a* coding sequence was replaced with a sequence encoding a *lexA::VP16* driver [88]. (B-D) MD assays of flies expressing the *fru-GAL4* driver together with (B) *Gr5a-RNAi* (C) *Gr43a-RNAi* (D) *Gr64f-RNAi*. (E) Male foreleg (upper panels), midleg (middle panels) and hindleg (bottom panels) of flies expressing *Gr5a-lexA* and *Gr64f-GAL4* drivers together with *lexAOP-tdTomato* and *UAS-Stinger* were imaged live under a fluorescent microscope. Yellow arrows indicate *Gr5a-*positive neurons and *Gr64f*-positive neurons. Scale bars represent 50 μm. (F) Male foreleg of flies expressing *Gr5a-lexA* and *Gr43a-GAL4* drivers together with *lexAOP-tdTomato* and *UAS-Stinger* were imaged live under a fluorescent microscope. (G) Male proboscis of flies expressing *Gr5a-lexA* and *Gr43a-GAL4* drivers together with *lexAOP-tdTomato* and *UAS-Stinger* were imaged live under a fluorescent microscope. Yellow arrows indicate *Gr5a-*positive neurons and *Gr43a*-positive neurons. Scale bars represent 50 μm. (H-I) MD assays of (H) *Gr64f-* and (D) *Gr43a-GAL4* drivers for the inactivation of synaptic transmission of *fru*-specific neurons among each *GAL4*-labelled neuron *via UAS>stop>TNT_active_; fru^FLP^*. Tested gustatory sugar receptors were selected based on a previous study [25].

By using the genetic intersectional method, we found that *Gr5a* is co-expressed with *Gr64f* in 5T - 1T of the male foreleg and 5T - 4T in midleg (Fig. 6E). However, *Gr5a* is co-expressed with *Gr64f* in 5T - 4T in the female foreleg/midleg and 5T in the female hindleg (Fig. S6H). In contrast, there are no *Gr5a*-positive cells expressing the fructose sensor Gr43a [26] in the male foreleg (Fig. 6F). Although no cells co-expressed Gr5a and Gr43a in the leg, several cells co-expressed Gr5a and Gr43a in the male proboscis (Fig. 6G). We were unable to detect any *fru*-positive cells expressing Gr64f in the male proboscis (Fig. S6I). When we expressed *UAS-TNT* only in male-specific *Gr64f*-positive neurons, we found that SMD was abolished; however, SMD remained intact in *Gr43a*-positive neurons (Fig. 6H-I). Gr proteins are known to function as heterodimeric or multimeric complexes [27–29]. In addition, Gr64f is required broadly as a co-receptor for the detection of sugars and works together with Gr5a protein to illicit behavioral responses to trehalose [30]. Collectively, these data suggest that co-expression of the sugar receptor Gr5a and its co-receptor Gr64f in male-specific leg sensory neurons is crucial for the sensory inputs underlying SMD behavior.

Next, we tested the role of pheromone processing molecules in male legs in the generation of SMD behavior [31]. The knockdown of LUSH, an odorant-binding protein [32] in *Gr5a*-positive neurons, but not in *Gr66a*-positive neurons, led to the abolishment of SMD behavior (Fig. 7A-B). SNMP1 is a member of the CD36-related protein family and functions as an important player for the rapid kinetics of pheromonal response in insects [33, 34]. We found that the expression of Snmp1 on the *snmp1* mutant background *via* the *Gr5a-GAL4* driver, but not the *Gr66a-GAL4* driver, could rescue SMD behavior (Fig. 7C-H), thus suggesting that the expression of the pheromone sensing proteins LUSH and Snmp1 in *Gr5a*-positive gustatory neurons is critical for generating SMD behavior. By using the genetic intersectional method, we found that the male antenna contains an abundance of *Snmp1*-positive cells but did not find any *Gr5a*-positive or ty*Snmp1*-positive cells (Fig. 7I). Surprisingly, we found one cell that was both *Snmp1*-positive and Gr5a-positive in the 2T of the male tarsus (Fig. 7J). Collectively, these data suggest that the expression of LUSH and SNMP1 in the male leg is crucial for sensory inputs for SMD behavior.

**Fig. 7.**
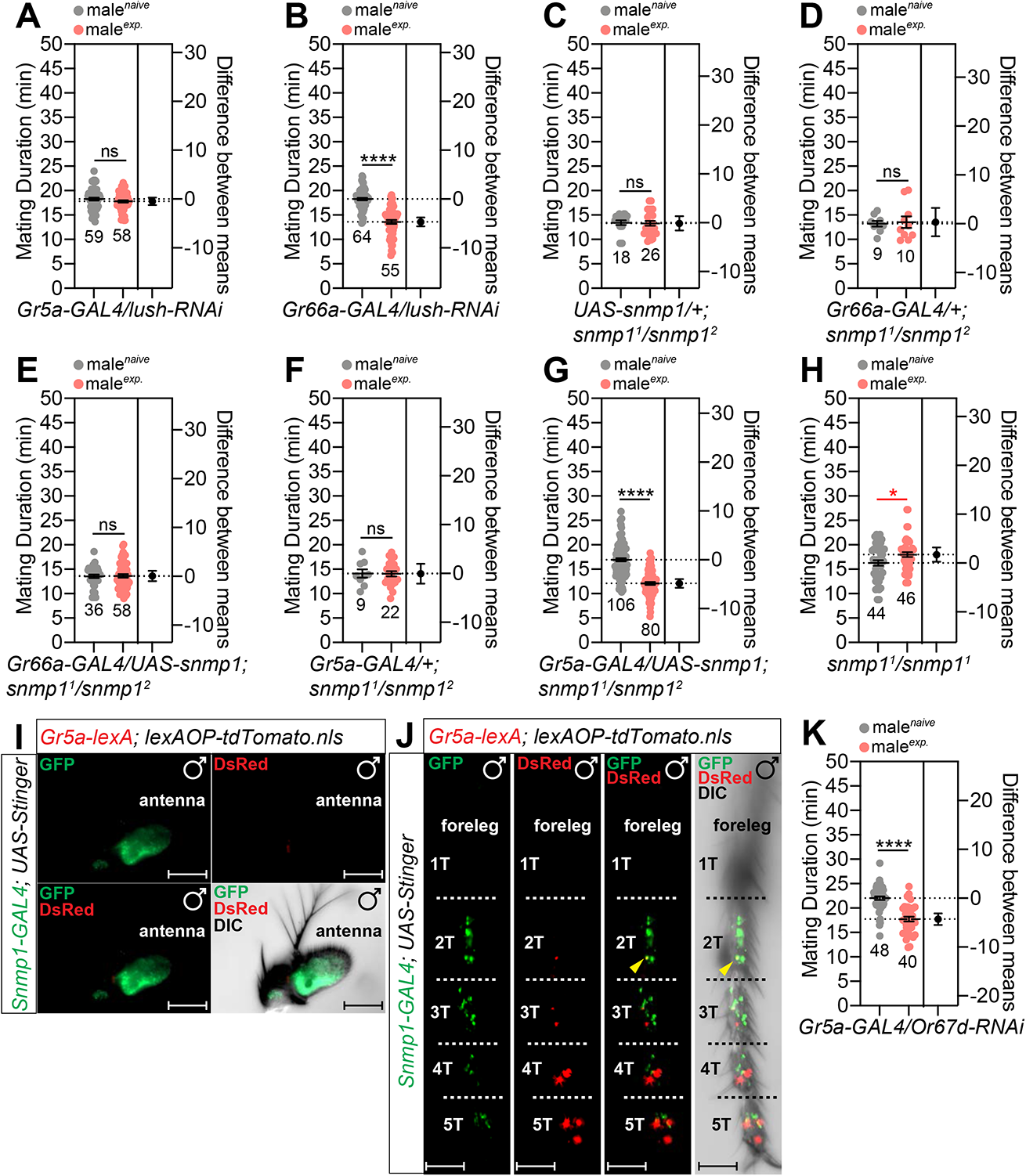
*OBPs* and *snmp1* are involved in detecting the sensory inputs for SMD behavior. (A-B) MD assays for *GAL4* mediated knockdown of LUSH *via UAS-lush-IR; UAS-dicer* (*lush-RNAi*) using (A) *Gr5a-GAL4* and (B) *Gr66a-GAL4* drivers. (C-H) MD assays of *snmp1* genetic rescue experiments. Genotypes are indicated below each graph. (I) Male antenna of flies expressing *Gr5a-lexA* and *Snmp1-GAL4* drivers together with *lexAOP-tdTomato* and *UAS-Stinger* were imaged live under a fluorescent microscope. Scale bars represent 50 μm. (J) Male foreleg of flies expressing *Gr5a-lexA* and *Snmp1-GAL4* drivers together with *lexAOP-tdTomato* and *UAS-Stinger* were imaged live under a fluorescent microscope. Yellow arrows indicate *Gr5a-*positive and *Snmp1*-positive neurons. Scale bars represent 50 μm. (K) MD assays for *GAL4* mediated knockdown of *Or67d via Or67d-RNAi* using *Gr5a-GAL4*.

Or67d is known to act as the receptor for male pheromone 11-*cis* vaccenyl acetate (cVA) [33, 35] and works together with SNMP1 for cVA-mediated olfactory responses [34]. We found that the knockdown of Or67d in Gr5a-positive or Gr66a-positive cells did not affect SMD (Fig. 7K, Fig. S7E-F). We found that the post-synaptic terminals of *Or67d-GAL4*-labeled neurons (Fig. S7G) and those of male-specific Or67d-GAL4-labeled neurons (Fig. S7H) target the antenna lobe in the male brain. These data suggest that OR67d-expressing olfactory neurons are not related to GR5a-expressing gustatory neurons in terms of providing sensory inputs for SMD behavior although OR67d requires SNMP1 function to respond to cVA. We also confirmed that Or67-GAL4 labels sensory neurons in the male antenna but not in the proboscis or foreleg (Fig. S7I). Collectively, these data suggest that Or67d and its ligand pheromone cVA are dispensable for SMD behavior *via* gustatory sensory inputs in males.

Next, we tested the importance of degenerin/epithelia Na^+^ channels (DEG/ENaC), *ppk23, ppk25* and *ppk29* in the excitability of pheromone-sensing cells [36, 37]. By using RNAi-mediated knockdown experiments, we found that *ppk25* and *ppk29*, but not *ppk23*, are crucial for generating SMD behavior in *Gr5a*-positive cells but not *Gr66a*-positive cells (Fig. 8A-F, Fig. S8A-C). By using the genetic intersectional method, we found that *ppk25* was co-expressed with *Gr5a* in 5T of the male foreleg and 4T of the midleg (Fig. 8G). We also found that *ppk29* was co-expressed with *Gr5a* in 2T and 4T of the male foreleg (Fig. 8H). However, we did not detect any cells that co-expressed *ppk23* and *Gr5a* in the legs of males (Fig. 8D). Of the Deg/ENaC sodium channel family, *ppk28* is reported to be expressed in gustatory neurons and is known to mediate the detection of water taste [38]. By using RNAi-mediated knockdown experiments, we found that *ppk28* is dispensable for SMD behavior in *Gr5a*-positive neurons (Fig. S8H-J). These data suggest that *ppk25*/*ppk29*, but not *ppk23*/*ppk28*, are crucial for pheromonal detection in the induction of SMD behavior in Gr5a-positive leg neurons in males.

**Fig. 8.**
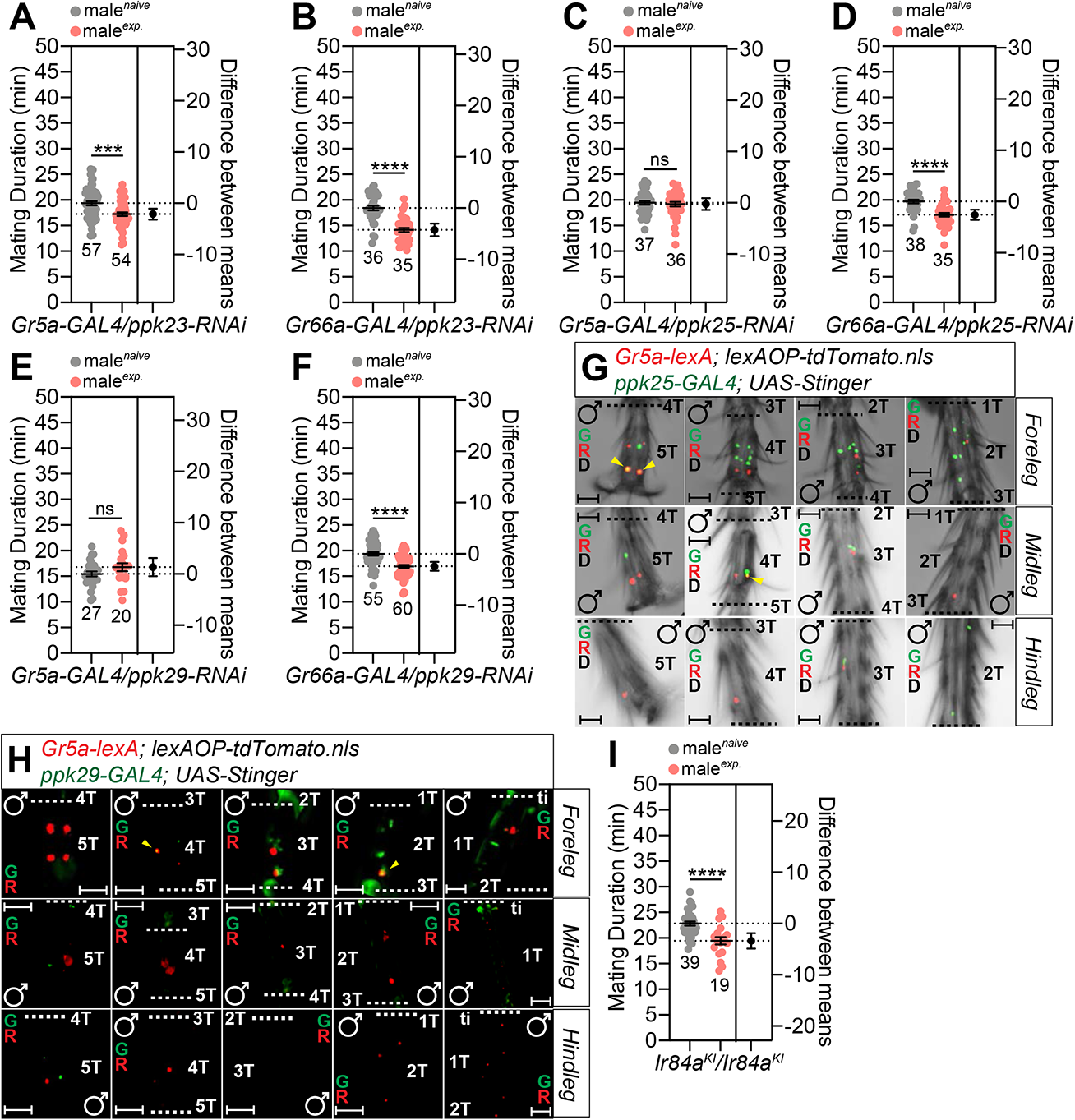
The DEG/ENaC channels *ppk25* and *ppk29* are crucial for detecting the sensory inputs for inducing SMD behavior. (A-B) MD assays for *GAL4* mediated knockdown of PPK23 *via ppk23-RNAi* using (A) *Gr5a-GAL4* and (B) *Gr66a-GAL4* drivers. (C-D) MD assays for *GAL4* mediated knockdown of PPK25 *via ppk25-RNAi* using (C) *Gr5a-GAL4* and (D) *Gr66a-GAL4* drivers. (E-F) MD assays for *GAL4* mediated knockdown of PPK29 *via ppk29-RNAi* using (E) *Gr5a-GAL4* and (F) *Gr66a-GAL4* drivers. (G) Male foreleg (upper panels), midleg (middle panels) and hindleg (bottom panels) of flies expressing *Gr5a-lexA* and *ppk25-GAL4* drivers together with *lexAOP-tdTomato* and *UAS-Stinger* were imaged live under a fluorescent microscope. Yellow arrows indicate *Gr5a-*positive and *ppk25*-positive neurons. Scale bars represent 50 μm. (H) Male foreleg (upper panels), midleg (middle panels) and hindleg (bottom panels) of flies expressing *Gr5a-lexA* and *ppk29-GAL4* drivers together with *lexAOP-tdTomato* and *UAS-Stinger* were imaged live under a fluorescent microscope. Yellow arrows indicate *Gr5a-*positive neurons and *Gr64f*-positive neurons. Scale bars represent 50 μm. (I) MD assay of *Ir84a-GAL4* knock-in null allele males of which the *Ir84a* coding sequence had been replaced with a sequence encoding a *GAL4* driver [89].

Ir84a is an olfactory receptor that detects phenylacetic acid (PAA) and phenylacetaldehyde (PA); these chemicals are present in various sources of fly food and enhance male courtship [39, 40]. PAA and PA act as a signal amplifier for olfactory responses by the DEG/ENaC channel, specifically *ppk25* [41, 42], and also activate the *fru^M^* mating circuitry [43]. SMD behavior was intact in *Ir84a* knockout flies which exhibited defects in the integration of signals arising from food and sex (Fig. 8I). By applying the RNAi-mediated pan-neuronal knockdown of Ir84a, we also found that the neuronal expression of the *Ir84a* gene is dispensable for SMD behavior (Fig. S8K-L). These data suggest that the *Ir84a*-mediated olfactory signaling pathway is distinct from the input pathways for SMD behavior.

Three *ppk* family members (*ppk23*, *ppk25* and *ppk29*) can sense the female pheromone 7,11-heptacosadiene [37] and express *fruitless*, a factor that is crucial for mating behavior in males [36]. By using RNAi-mediated knockdown, we found that the expression of *ppk25*/*ppk29* in *fru*-positive cells is crucial for SMD behavior, but not *ppk23* expression (Fig. 9A-C). By using the genetic intersectional method, we identified that *ppk23* was co-expressed with *fru* in 5T - 2T of the male foreleg and 2T of the hindleg (Fig. 9D). We also found that *ppk25* was co-expressed with *fru* in 5T - 2T of the male foreleg and 4T of the midleg (Fig. 9E) and that *ppk29* was co-expressed with *fru* in 5T - 2T of the male foreleg (Fig. 9F, Fig. S9A). We also confirmed that *ppk29-GAL4* labels cells only in males and not in females (Fig. S9B-C). These data suggest that the expression of *ppk25* and *ppk29* in *fru*-positive male-specific cells is crucial for SMD behavior.

**Fig. 9.**
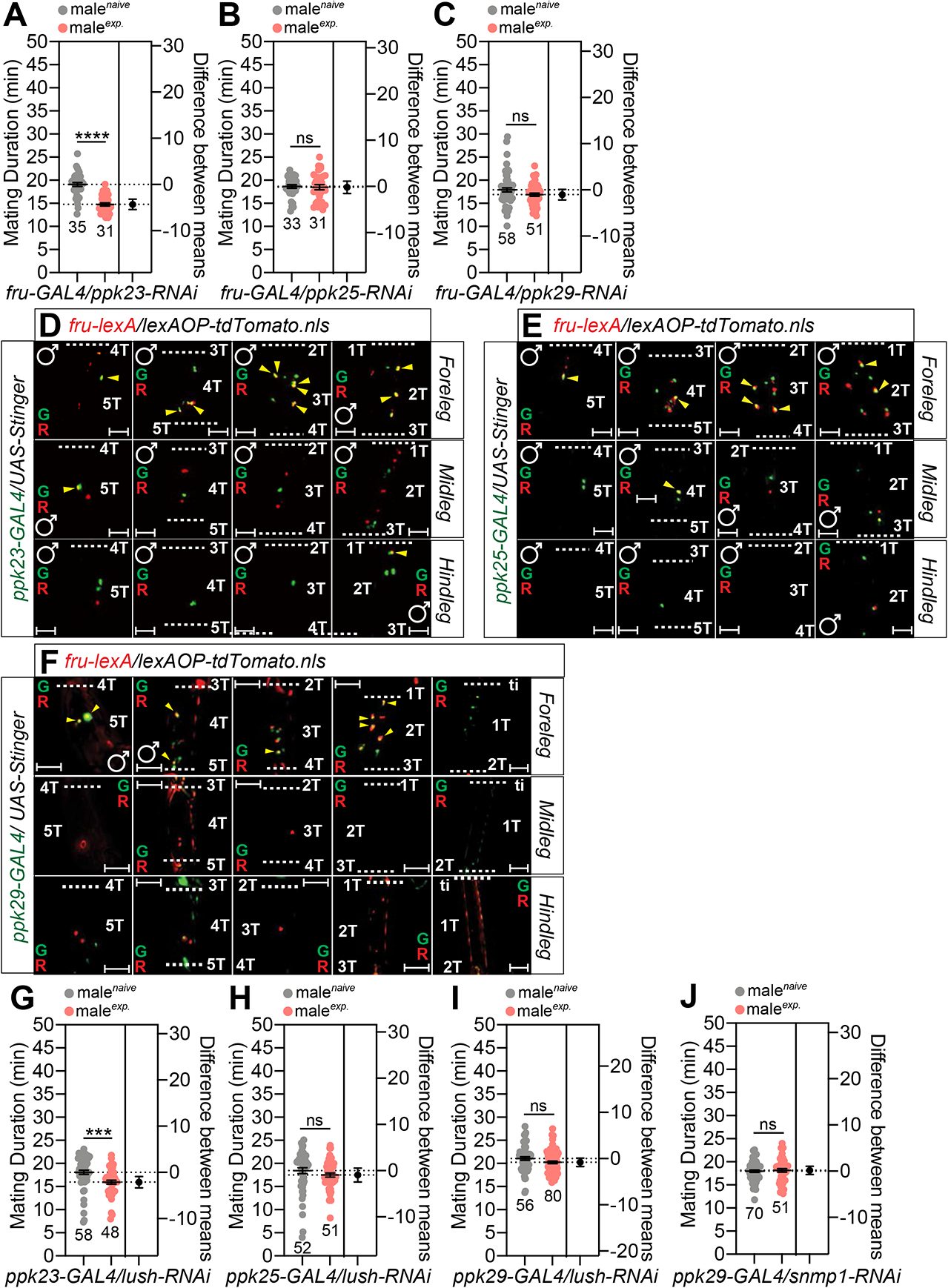
The expression of PPK25 and PPK29 in *fru*-positive sexually dimorphic cells is crucial for detecting the sensory inputs for inducing SMD behavior. (A-C) MD assays for *GAL4* mediated knockdown of (A) PPK23, (B) PPK25, and (C) PPK29 *via ppk23-RNAi*, *ppk25-RNAi*, and *ppk29-RNAi* using the *fru-GAL4* driver. (D-F) Male foreleg (upper panels), midleg (middle panels) and hindleg (bottom panels) of flies expressing *fru-lexA* and (D) *ppk23-GAL4*, (E) *ppk25-GAL4*, and (F) *ppk29-GAL4* drivers together with *lexAOP-tdTomato* and *UAS-Stinger* were imaged live under a fluorescent microscope. Yellow arrows indicate *fru-*positive and *ppk23-*, *ppk25-*, or *ppk29*-positive neurons. Scale bars represent 50 μm. (G-I) MD assays for *GAL4* mediated knockdown of LUSH *via lush-RNAi* using (G) *ppk23-GAL4*, (H) *ppk25-GAL4*, (I) *ppk29-GAL4* drivers. (J) MD assays for *GAL4* mediated knockdown of SNMP1 *via snmp1-RNAi* using the *ppk29-GAL4* driver

Next, to decipher whether DEG/NaC channel-expressing pheromone sensing neurons require the function of OBP, we expressed *lush-RNAi* using *ppk23*-, *ppk25*-and *ppk29-GAL4* drivers to knockdown LUSH in each channel-expressing neuron. The knockdown of LUSH in *ppk25*- and *ppk29-GAL4* labeled cells, but not in *ppk23-GAL4* labeled cells, led to a disturbance in SMD behavior, thus suggesting that LUSH functions in *ppk25*-and *ppk29*-positive neurons to detect pheromones and elicit SMD behavior (Fig. 9G-I). The knockdown of SNMP1 in *ppk29-GAL4-*labeled neurons also inhibited SMD behavior (Fig. 9J), thus suggesting that SNMP1 also functions in *ppk29*-positive neurons to induce SMD behavior.

The *Drosophila melanogaster* genome bears two members of the SNMP/CD36 gene family; the proteins these genes encode are expressed in distinct cells [44, 45]. SNMP2 is known to contribute to gender recognition during courtship; however, its precise functional role remains unknown [45, 46]. To compare the function of SNMP2 with SNMP1, a factor that is crucial for SMD behavior, we reduced the gene expression of SNMP2 in *ppk23*-, *ppk25-*, *ppk29-GAL4* expressing pheromone sensing neurons and found that SNMP2 is dispensable in these pheromone-sensing neurons for eliciting SMD behavior (Fig. S9D-F). We also found that SNMP2 was not required for SMD behavior in *Gr5a*- and *Gr64f-GAL4* labeled sugar sensing neurons (Fig. S9G-H). These data suggest that SNMP1, but not SNMP2, is specifically involved in pheromone detection for SMD behavior in the male leg system.

The legs of insects are associated with multiple mechanoreceptor types nearby and each mechanoreceptor type is sensitive to a particular range of mechanical stimuli [47]. To decipher the involvement of mechanosensory inputs in generating SMD behavior, we screened the functionally characterized channels involved in mechanosensation using a *Gr5a-GAL4* driver [48–52]. Of the thirteen receptors screened by RNAi knock-down, we only found that the knock-down of *nanchung* and *pyrexia* in *Gr5a*-positive cells affected SMD behavior (Fig. 10A-M, Fig. S10A-M), thus suggesting that *nanchung* and *pyrexia* specifically modulate the mechanosensory inputs in *Gr5a*-positive cells but not fructose receptor *Gr43a*- or *Gr66a*-positive (salt/avoidance cells) cells to generate SMD behavior (Fig. 10N-P).

**Fig. 10.**
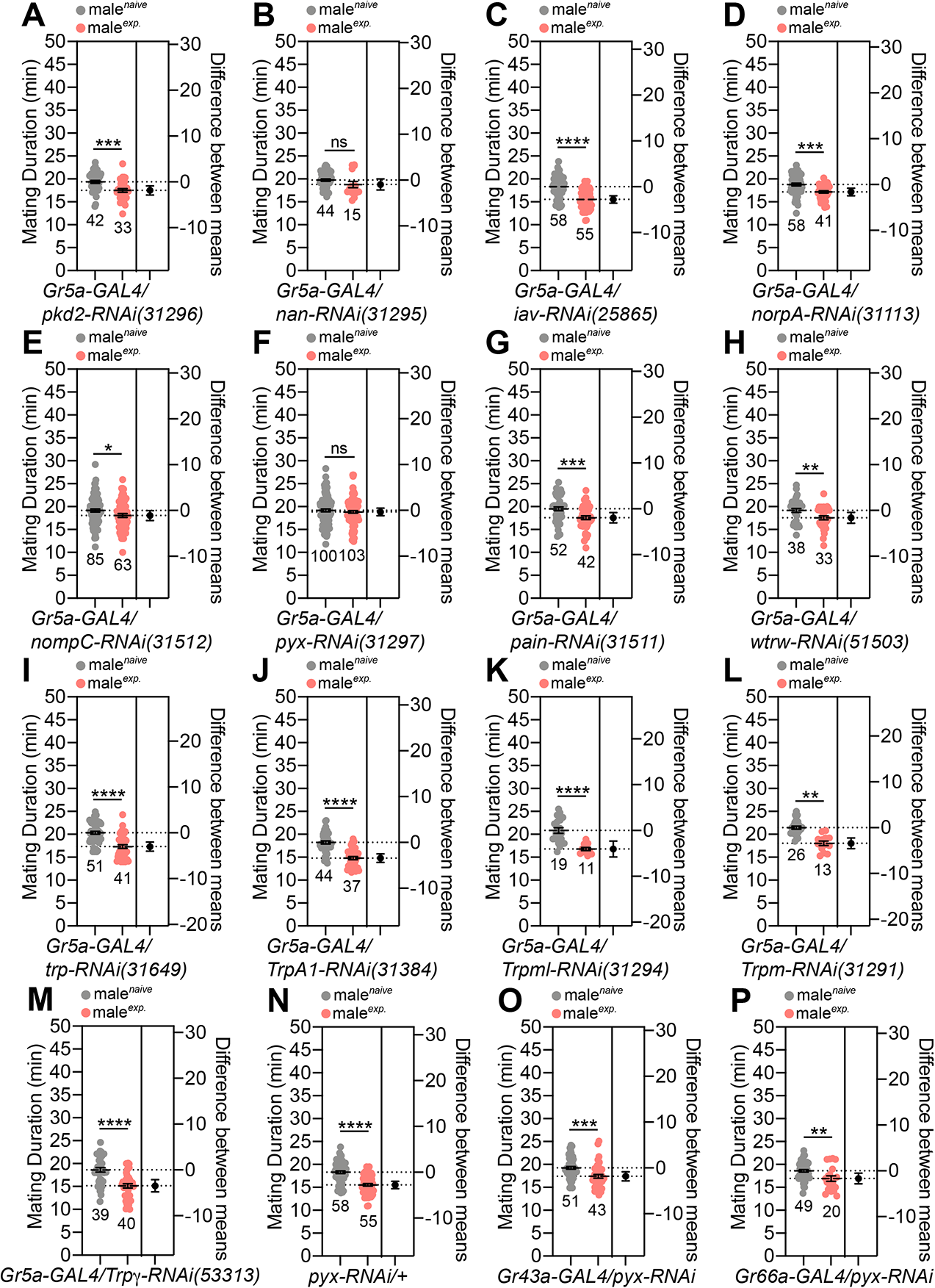
Specific channels for auditory and mechanosensory sensing are associated with SMD behavior. (A-M) MD assays for *GAL4* mediated knockdown of channels and receptors mediate auditory, mechanosensory and thermal sensing using the *Gr5a-GAL4* driver. Tested mechanosensation-related receptors were selected based on previous studies [49,51,52]. (N) Control experiments for MD assays in Fig. 10F **and 10O-P**. (O-P) MD assays for *GAL4* mediated knockdown of PYX via *pyx-RNAi* using (O) *Gr43a-GAL4* and (P) *Gr66a-GAL4* drivers.

By using the genetic intersectional method, we found a single cell expressing *Gr5a-lexA* together with *pyx-GAL4* in the 4T of the male foreleg (Fig. 11A); however, we could not find any of these relevant cells in the female foreleg (Fig. 11B). Next, we found that *pyx-GAL4* was expressed in approximately 10 cells in the male foreleg tarsus; of these, 8 were in the 1T (Fig. S11A). Interestingly, we found approximately 8 cells in the male proboscis which could be labeled by both *pyx-Gal4* and *Gr5a-Gal4*, thus suggesting that *pyx* might exert function in the gustatory system.

**Fig. 11.**
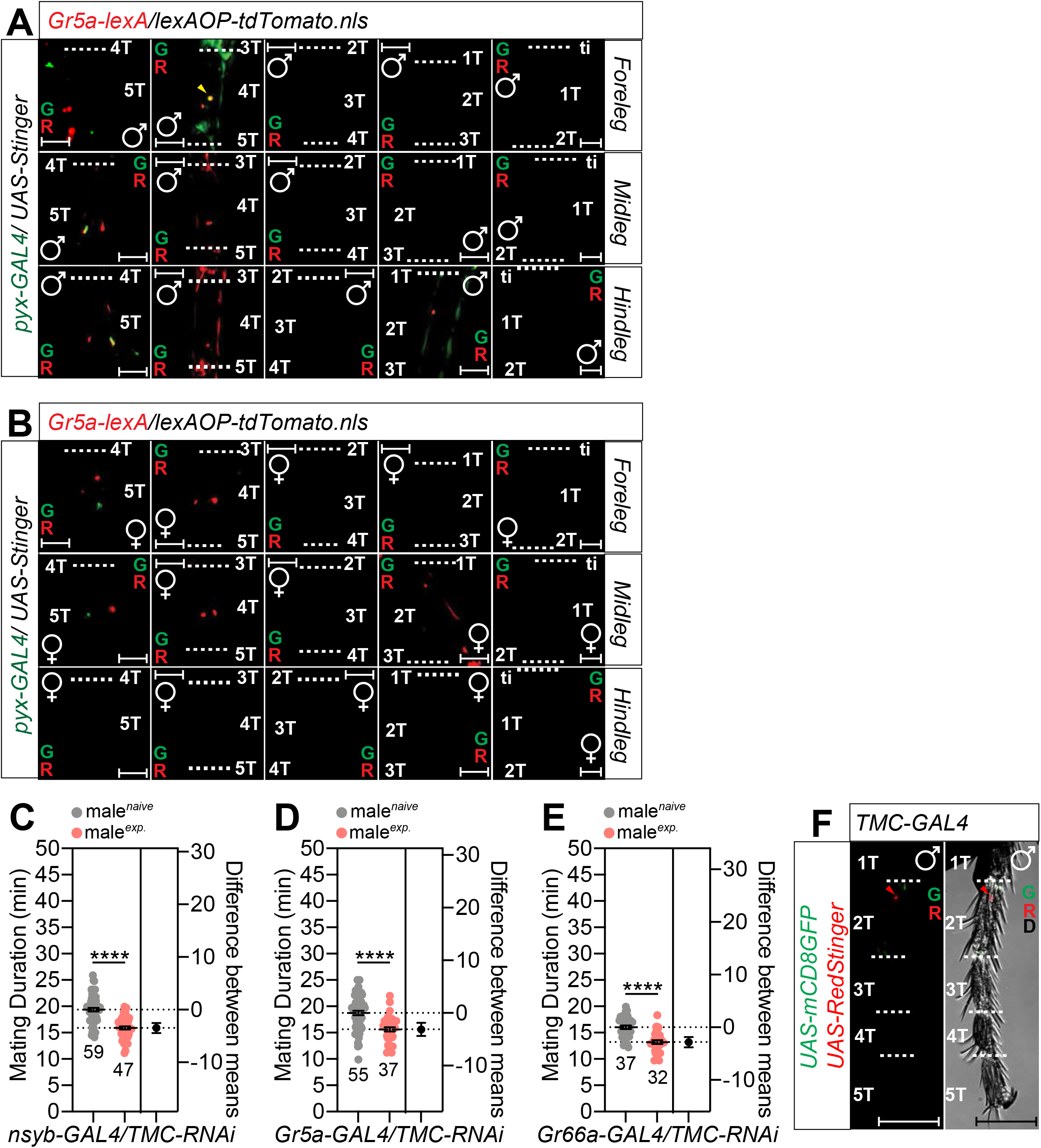
The expression of *pyx* in *Gr5a*-positive sexually dimorphic cells is crucial for detecting the sensory inputs for inducing SMD behavior. (A-B) (A) Male and (B) female foreleg (upper panels), midleg (middle panels), and hindleg (bottom panels) of flies expressing *Gr5a-lexA* and *pyx-GAL4* driver together with *lexAOP-tdTomato* and *UAS-Stinger* were imaged live under a fluorescent microscope. Yellow arrows indicate *Gr5a-*positive neurons and *pyx*-positive neurons. Scale bars represent 50 μm. (C-E) MD assays for *GAL4* mediated knockdown of TMC *via TMC-RNAi* using (C) *nSyb-GAL4*, (D) *Gr5a-GAL4*, and (E) *Gr66a-GAL4* drivers. (F) Male foreleg of flies expressing *TMC-GAL4* together with *UAS-RedStinger, UAS-mCD8GFP* were imaged live under a fluorescent microscope. Scale bars represent 50 μm.

The recently discovered evolutionarily conserved mechanosensitive channel, transmembrane channel-like (*TMC*) [53] can sense the hardness of substrates and can discriminate the textures of the foreleg contact region [54]. It has been reported that sweet neurons inhibit texture discrimination by signaling *TMC*-expressing mechanosensitive neurons in female *Drosophila* [54], thus suggesting that sugar perception and mechanosensation through *TMC* might be associated with the male foreleg. We reduced the expression of *TMC* by performing RNAi-mediated knockdown experiments and found that pan-neuronal (Fig. 11C), sugar cell (Fig. 11D), or bitter cell (Fig. 11E) expression of *TMC* was not required with the sensory inputs for SMD behavior. We also found that *TMC-GAL4* rarely labelled cells in the male leg (Fig. 11J and Fig. S11C), thus suggesting that the texture discrimination function of *TMC* is not required for SMD behavior.

We summarized the identified cells that expressed *fru, Gr5a, Gr64f, ppk25, ppk29, snmp1*, and *pyx* in the male or female legs as diagrams (Fig. S11D). In summary, we identified the male-specific sensory cells that integrate multisensory inputs from the sugar receptors (*Gr5a* and *Gr64f*), pheromone sensing machinery OBP and CD36 family (*lush, snmp1*), pheromone receptors (*ppk25* and *ppk29*), auditory/mechanosensory receptor (*nan*), and transient receptor potential channel (*pyx*) to generate input signals for SMD.

To determine whether the temporal activation of *Gr5a*-positive neurons may generate SMD behavior in the absence of sexual experiences, we expressed the heat-sensitive *Drosophila* cation channel *TrpA1* in *Gr5a*-positive cells and then transferred the experimental group only to the activation temperature (29℃). Surprisingly, the flies expressing *TrpA1* in *Gr5a*-positive neurons at the activation temperature showed a shorter mating duration than those that remained at 22°C (Fig. 12A). Neither the genetic control (Fig. 12B and C) nor the flies expressing *shi^ts^* that could disrupt synaptic transmission in a temperature-sensitive fashion (Fig. 12E) showed changes in their mating duration between 22°C and 29°C. These findings indicate that the stimulation of *Gr5a*-positive neurons is sufficient to generate SMD behavior.

**Fig. 12.**
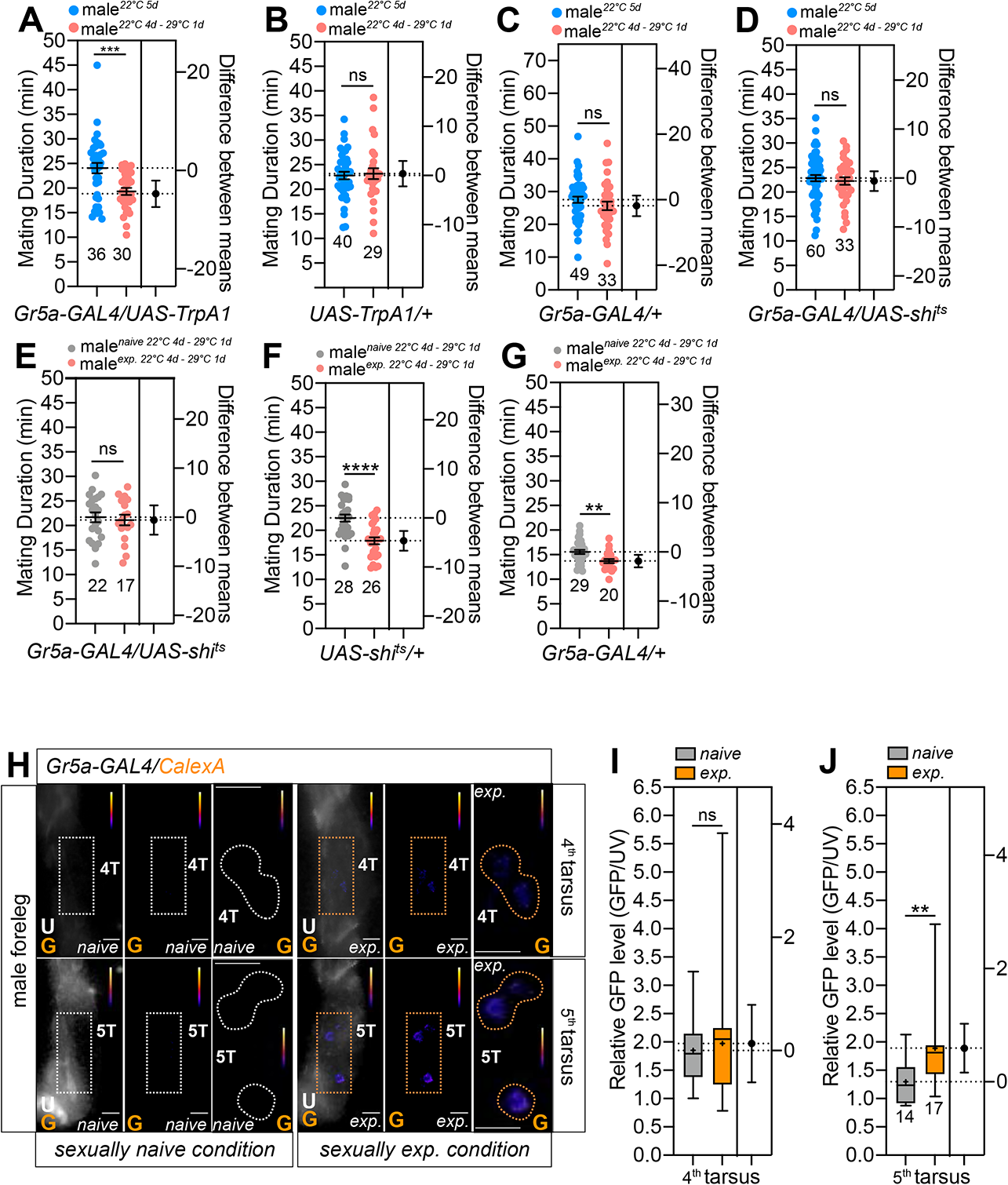
Temporal activation of Gr5a neurons induced SMD behavior without sexual experiences (A-D) MD assay for the temporal temperature-shift of flies expressing *UAS-TrpA1* or *UAS-shi^ts^* by *Gr5a-GAL4*. Genotypes are labelled below the graph. Blue groups were reared at 22°C for five days and red groups were reared at 22°C for four days and moved to 29°C overnight. (E-G) MD assay for the temporal temperature-shift of flies expressing *UAS-shi^ts^* by *Gr5a-GAL4*. Genotypes are labelled below the graph. Grey groups reared at 22°C for four days and moved to 29°C for overnight. Red groups reared at 22°C for four days and moved to 29°C overnight with sexual experiences. (H) Different levels of neural activity of the 4^th^ and 5^th^ sensory neurons as revealed by the CaLexA system in naive *versus* mated male flies. Male flies expressing *Gr5a-GAL4* along with *LexAop-CD2-GFP, UAS-mLexA-VP16-NFAT and LexAop-CD8-GFP-2A-CD8-GFP* were dissected after at least 10 days of growth (mated male flies had 1-day of sexual experience with virgin females). GFP is pseudo-colored as “red hot”. Dashed boxes represent the magnified area of interest and show the right section of each condition. Dashed circles represent the location of *Gr5a*-positive cells. White colors represent the naïve condition while the yellow color represents the experienced condition. Scale bars represent 20 µm. (I and J) Quantification of GFP fluorescence. GFP fluorescence of the 4^th^ (I) or 5^th^ (J) tarsus was normalized to that in auto-fluorescence. The conditions of flies are described above: naïve, naïve male flies; exp., male flies with sexual experience. Bars represent the mean of the normalized GFP fluorescence level with error bars representing the SEM. Asterisks represent significant differences, as revealed by the Student’s t test and ns represents non-significant difference (*p < 0.05, **p < 0.01, ***p < 0.001).

By using the expression of *shi^ts^* with *Gr5a-GAL4*, we then attempted to inhibit the synaptic transmission of *Gr5a*-positive neurons during sexual experiences. We discovered that inhibiting *Gr5a*-positive neurons during sexual interactions by increasing the temperature to 29°C could impair SMD behavior (Fig. 12E). The genetic control exhibited no such result (Fig. 12F and G). These findings imply that the neural stimulation of *Gr5a*-positive neurons is a crucial trigger for SMD behavior.

To determine whether neuronal activities undergo alterations in neurons associated with SMD, we utilized the CaLexA (calcium-dependent nuclear import of *LexA*) system [55]. This system is based on the activity-dependent nuclear import of the transcription factor nuclear factor of activated T cells (*NFAT*). Because SMD needs at least 6–12 h of sexual interaction, repeated sensory inputs might theoretically lead to the buildup of the modified transcription factor within the nucleus of activated neurons *in vivo*. Indeed, sexual encounters affected the neural activity of some *Gr5a-GAL4*-labeled neurons. Male flies with sexual experience and carrying *Gr5a-GAL4* and *LexAop-CD2-GFP; UAS-mLexA-VP16-NFAT, LexAop-CD8-GFP-2A-CD8-GFP* exhibited strong fluorescence in the 5^th^ tarsus following an overnight sexual experience. In contrast, no similar signals were identified in males with no prior experience. In contrast to *Gr5a*-positive neurons in the 5th tarsus, cells in the 4^th^ tarsus did not exhibit a significant increase in GFP fluorescence (Fig. 12H-J), thus indicating that sexual encounters change the neuronal activity of *Gr5a* cells in the 5^th^ tarsus.

To explore the adaptive value of SMD, we developed a theoretical model to test the adaptive value of SMD behavior based on the marginal value theorem [56, 57] (Box. S1). This model assumes that both the benefits and costs of mate guarding accumulate over time, but with different aspects. The benefit refers to the number of eggs fertilized by the guarding male while the costs refer to the guarding-associated potential costs such as increased predation risk or the loss of opportunities for other forms of mating or foraging activity [58]. The model suggests that shortened mating durations can be preferred in experienced males if (1) experienced males can fertilize a fewer number of eggs in total than naïve males (Fig. 13A) and that the rate of fertilization is (2) faster (Fig. 13B) or (3) slower (Fig. 13C) for experienced males while the total number of eggs that can be fertilized remains the same as for naïve males, and/or 4) the costs accumulate faster in experienced males (Fig. 13D). Next, we empirically tested which scenario(s) could explain the observed SMD behavior. We focused on testing scenarios 1-3 but not 4, firstly because it was hard to identify a rationale for how the costs of mate guarding differ between experienced and naïve males and secondly, to experimentally manipulate the costs.

**Fig. 13.**
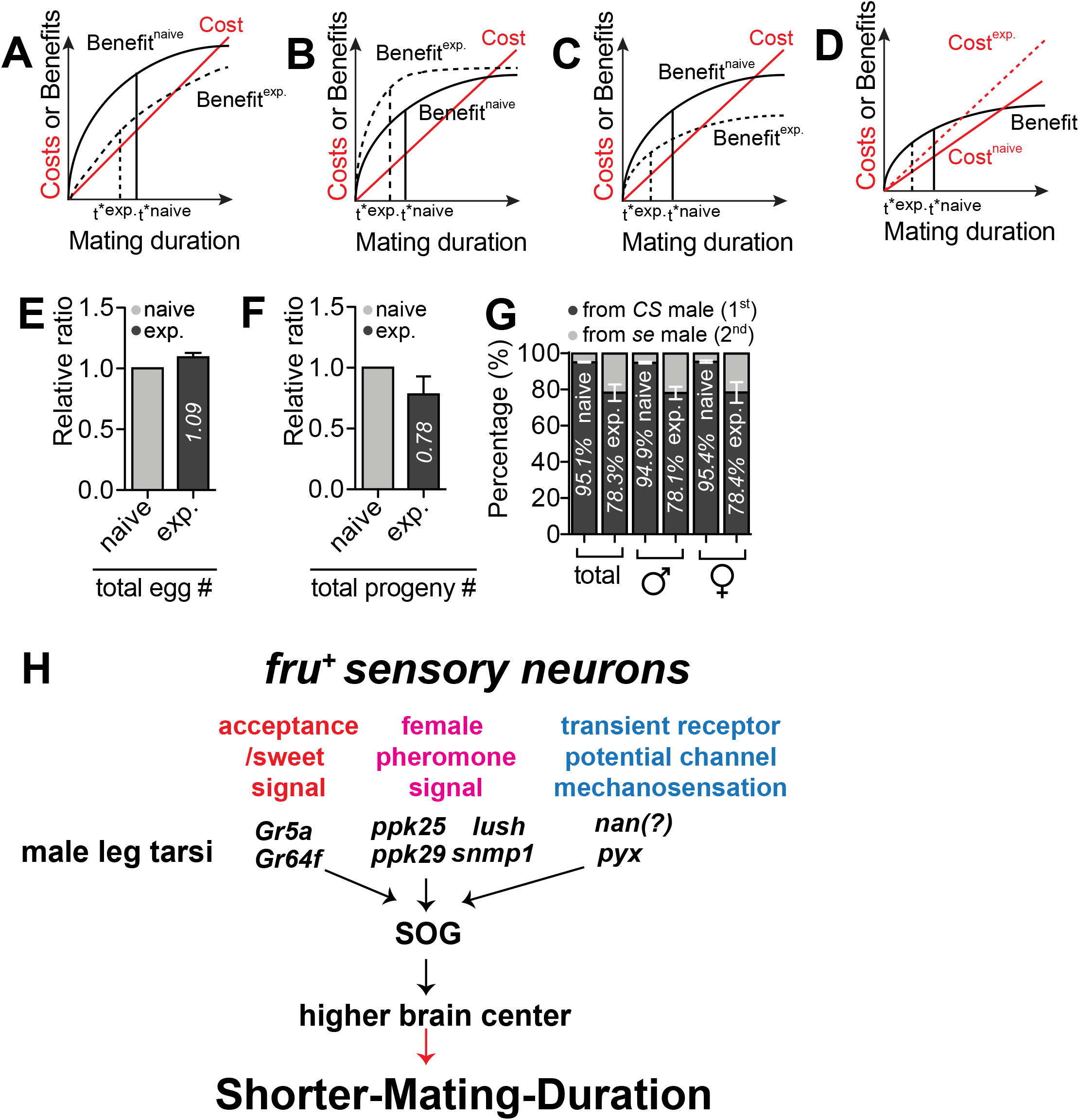
Adaptive benefits of SMD behavior. (A)-(D) show the four different scenarios by which SMD can evolve (see **Box. S1**). SMD can evolve when α gets larger (A), β gets smaller (B), γ gets larger (assuming β/α < γ < e*β/α) (C) and/or γ gets smaller (assuming γ > e*β/α) (D). (E) Relative ratio of total egg number comparing the eggs produced by females mated with naïve males to the eggs produced by females that mated with experienced males. (naïve = control bar for comparing, exp. = eggs from naïve males/eggs from exp. males). (F) Relative ratio of total progeny number comparing the progeny produced by the females mated with naïve males to the progeny produced by females mated with experienced males. (naïve = control bar for comparing, exp. = progeny from naïve males/progeny from exp. males). (G) Percentage of progeny originated from *sepia* (*se*) male *versus CS* male. *se* male was introduced to *se* female as first mate then followed by *CS* males as the second mate. The eye color of progeny was counted and interpreted as the source of the father; for detailed methods, see the **EXPERIMENTAL PROCEDURES**. (H) Summary of this study showing the multisensory inputs modulating SMD behavior.

We found that the total number of eggs produced by females that mated with experienced males was comparable to those that mated with naïve males (Fig. 13E); however, the number of progeny from the experienced males was significantly lower than those from naïve males (Fig. 13F). When females that mated with an experienced or naïve male were subsequently introduced to another male after a certain time period, the number of progenies arising from the experienced males was also significantly fewer than those from the naïve males (Fig. 13G and Fig. S12A). This suggests that (i) the number of sperm from experienced males for fertilization in a given period of time was lower than that from naïve males or ii) females reduced the use of sperm from experienced males for fertilization when they had a choice. These results support scenario 1 and potentially scenario 3 in that SMD has evolved because the reproductive payoffs of experienced males through mate-guarding are consistently lower than those of naïve males.

## DISCUSSION

Our study provides new lines of evidence that male flies invest less time for mating duration when they are sexually experienced. Males retain a memory of sexual experience for several hours and economize mating duration accordingly (Fig. 1). This behavior relies primarily on gustatory input from the male forelegs, indicating that contact chemoreception is required for SMD induction (Fig. 2 and Fig. 3). Sugar cells expressing *Gr5a*, but not bitter cells expressing *Gr66a*, were found to be involved in the induction of SMD (Fig. 4). We also found that male-specific, *fru-*expressing *Gr5a*-positive sensory neurons are required to recognize the presence of females (Fig. 5). Sugar receptors such as *Gr5a/Gr64f*, but not fructose sensor *Gr43a*, are important for the sensory inputs required for SMD behavior (Fig. 6). Chemosensory proteins such as *lush* and *SNMP1*, as well as female pheromone receptors (DEG/ENaC channel *ppk25* and *ppk29*) are important for generating SMD (Fig. 7, Fig 8, and Fig.9). Of the screened auditory/mechanosensory receptors and transient receptor potential channels, *nanchung* and *pyrexia* were found to be crucial in the generation of SMD in sugar cells (Fig. 10 and Fig. 11). Using both theoretical and empirical approaches, we further showed that SMD represents the adaptive behavioral plasticity of male flies (Fig. 13).

Our earlier study found that previous exposure to rivals lengthens mating duration, a behavior that is referred to as longer-mating-duration (LMD) [14, 17]. The two behavioral circuits for LMD and SMD might have evolved independently since they use different sensory cues for detecting ‘rivals’ or ‘females’ for ‘sexual competition’ or ‘mating investment’, respectively. We propose that multisensory inputs from male forelegs detect the chemical signals from the female body and contribute to the determination of mating investment in male *Drosophila melanogaster* (Fig. 13H). The visual inputs from the male’s compound eye are the most crucial sensory cue to generate LMD [17]; however, multisensory inputs from the foreleg are required to induce SMD (Fig. 13H). To confirm that LMD does not require female pheromone signaling, we reduced the expression of the female pheromone receptor *ppk29* in all neuronal populations using RNAi-mediated knockdown experiments and found that the neuronal expression of *ppk29* is only essential for SMD but not for LMD behavior (Fig. S12B-C). Consistent with our previous report on the different neural circuitry for LMD and SMD [59], these data clearly show that male flies use different sensory modalities to generate LMD or SMD, respectively.

In our sugar receptor screening for SMD behavior, we found that only *Gr5a* and *Gr64f* were required for SMD behavior (Fig. 6A, B, and D). The other known sugar receptors (*Gr61a*, *Gr64a*, *Gr64b*, *Gr64c*, *Gr64d*, and *Gr64e*) are not required for SMD behavior (Fig. S6A-G). Fujii et al reported the expression code for specific sweet neurons in labial palp and tarsal sensilla [25]. In this code, *Gr5a*- and *Gr64f*-positive but *Gr43a*-negative neurons are referred to as “f4b”, “f4s”, “f5s”, and “f5b”. In the foreleg, the hair cells expressing *Gr43a* do not express *Gr5a* [25]. Gr43a is the fructose sensor and is co-expressed with Gr61a[26]. In summary, we suggest that the sugar receptors *Gr5a* and *Gr64f* in *fruitless*-positive cells provide crucial sensory information for SMD behavior.

It is known that the type of neurons expressing *ppk23*, *ppk25*, and *ppk29* is referred to as a “female” cell (F cell) from its responses to female aphrodisiac pheromones; the other type of neurons, expressing *ppk23 and ppk29* but not *ppk25*, is referred to as the “male” cell (“M” cell) from its response to male anti-aphrodisiac pheromones. M and F cells both express *fruitless* gene [36] SMD requires female pheromonal inputs through the contact chemoreception pathway in males (Fig. 2K-M). We also identified that there are *ppk25*- and/or *ppk29*-positive neurons among *Gr5a*-positive sugar detecting neurons (Fig. 8A-H). Thus, we hypothesize that F cells, which can detect sugar taste, are responsible for SMD behavior. Several groups have reported that *ppk23*-expressing cells respond to pheromones but not to water, salt, or sugars; in addition, this response is abolished by the mutation of either *ppk23* or *ppk29* [20,60,61]. Genetic rescue studies revealed that although all three subunits are co-expressed and function in the gustatory cells required for the activation of courtship by pheromones, each has a non-redundant function within these cells [62, 63]. Thus, we suggest that the *ppk25* and *ppk29* receptors expressed in *fruitless*-positive “F cells” are critical for detecting female body pheromones *via* contact chemoreception and generating SMD behavior.

One of the intriguing findings of this report is that *Gr5a*-positive taste neurons also express the female pheromone receptors *ppk25* and *ppk29*. To further validate our experimental data, we made use of a scRNA sequencing dataset of fruit flies that is available on the SCope website [64]. As described previously, the male fly’s front leg contains the sensory cells that can integrate gustatory, pheromonal, and mechanosensory inputs from sexual experiences. Using the annotation function of SCope, we confirmed that these sensory organs exhibited comparable cell population distribution patterns (Fig. S15A). In addition, the SCope annotation algorithm predicted that the majority of pheromone-sensing neurons are located on the wing among these three sensory organs (Fig. S15B). Using the comparison function of SCope, we discovered that males have a greater number of sensory neurons annotated both as ‘pheromone-sensing neurons’ and ‘gustatory receptor neurons’. Using the available categories of annotations in SCope that are associated with sensory information, we identified sensory cell populations that were annotated both as ‘gustatory receptor neurons’ and ‘pheromone sensing neurons’ as well as ‘mechanosensory neurons’ (Fig. S15C). The population of sensory cells we discovered exhibits sexual dimorphism (the far-right panel in Fig. S15C).

Numerous *Gr5a*-positive cells overlapped with the *ppk29*-expressing neurons that detect pheromones. In addition, we discovered that *ppk29*-positive cells overlap with mechanosensory neurons. One *nan*-positive cell was discovered within a pheromone-sensing neuron; this overlapped with *ppk29*-positive cells (Fig. S16A). When we focused on the mechanosensory neuron cluster of the leg chordotonal organ, we discovered that the majority of *nan*-positive cells were localized in this cluster. In this cluster, nan overlapped significantly with leg taste bristle chemosensory neurons and *ppk29*-positive neurons. Throughout this cluster, the *ppk25*-positive cells co-expressed nan and are classified as leg taste bristle chemosensory neurons. Intriguingly, this cluster of cells contained numerous cells expressing both *Gr5a* and *nan* (Fig. S16B).

Next, we investigated scRNA seq data from the legs. We were able to distinguish top, middle, and bottom clusters based on the three independent annotations of sensory neurons, gustatory receptor neurons, and mechanosensory neurons of the leg chordotonal organ. In the top cluster, we discovered a cell that co-expressed *Gr5a* and *ppk29*. In the middle cluster, cells expressing *ppk25* and *ppk29* were identified as gustatory receptor neurons. In the lowest cluster, cells expressing *ppk25* or *ppk29* were identified as mechanosensory neurons of the leg chordotonal organ (Fig. 14A). When we focused on male-specific *fru*-positive cells, the top cluster contained several *fru*-positive cells that expressed *ppk25* and *ppk29*. Numerous mid-panel cells were also identified as *fru*-positive gustatory receptor neurons; however, *ppk25*- and *ppk29*-positive male-specific cells were not detected in the mid panel. Many nan-positive cells were also *fru*-positive, as seen in the bottom panel of Fig. 14A. We also discovered numerous cells expressing *nan* and *ppk25*, *ppk25* and *fru*, or *fru* and *nan* together. One cell expressed these three marker genes, as shown in the bottom panel (Fig. 14B).

**Fig. 14.**
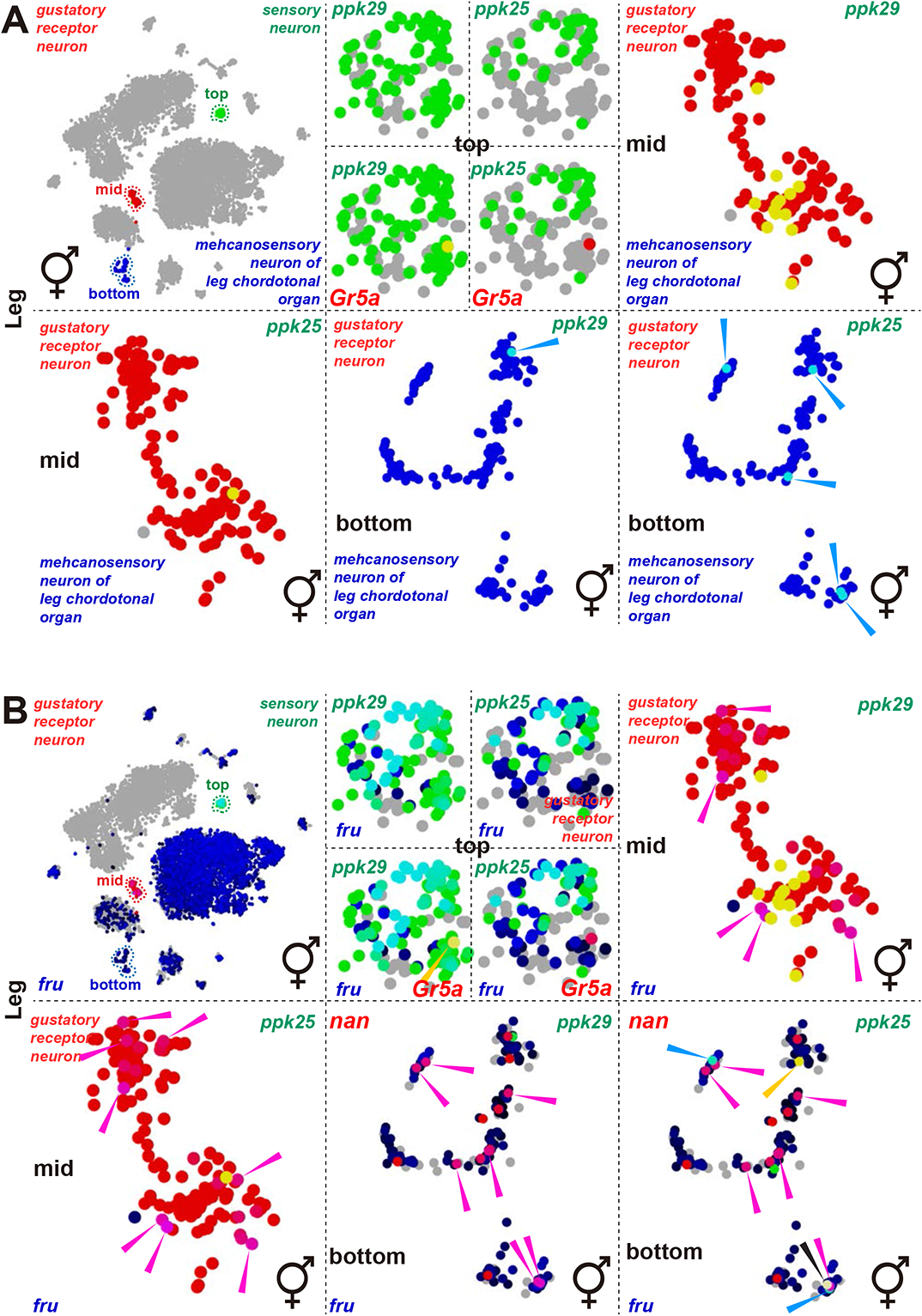
snRNAseq of adult fly legs [64] confirmed the expanded co-expression of the sensory receptor genes identified in this study. (A-B) Annotations and gene names are color-coded using red, green and blue words. When cells overlap, the color of the dots is either yellow, cyan or magenta. Colored arrows indicate cells that have been double- or triple-labeled by annotations and genes.

Next, we investigated the wing scRNA sequencing data since an examination of the whole tissue scRNA data suggested that the wing features the same sensory cell population that expresses SMD-essential genes. Using annotations such as gustatory receptor neurons, pheromone-sensing neurons, and mechano-sensory neurons from the top, middle, and bottom clusters, we observed clustering patterns that were comparable to the leg. Numerous pheromone-sensing neurons and a few gustatory receptor neurons co-expressed *Gr5a* and *ppk29* in these three clusters. An abundance of mechanosensory neurons expressed *ppk29* (Fig. S17A). In contrast to the leg, several cells designated as gustatory receptor neurons were *fru*-positive, thus indicating that many functions of gustatory sensing in the wing appear to be male-specific. Two cells in the pheromone-sensing neuron cluster of the wing expressed *Gr5a*, *ppk29*, and *fru*. Similar to the leg, the wing included tiny clusters of both *fru*-positive and *nan*-positive cells, thus indicating that *nan*-expressing male-specific cells are crucial for SMD and may be implicated in the mating behavior of other males. We also discovered a cluster of *ppk29*-positive but *ppk25*-negative pheromone-sensing neurons (Fig. S17B).

Next, we investigated the previously published FlyCellAtlas (FCA), a single-cell RNA-seq dataset [64], to determine if other organs exhibit the same gene expression profiles as the leg and the wing. We reviewed the expression levels of essential marker genes for SMD behavior in several sensory organs, including the leg, wing, proboscis, antenna, trachea, and oenocyte, and concluded that these genes are expressed comparably in the leg and wing, but not in other sensory organs (Fig. 15A-F). In addition, we discovered that *Gr5a* and *Gr64f* are expressed in gustatory receptor neurons and other sensory neurons in the leg, which are pheromone-sensing neurons in the wing, as we continued to divide cell types (Fig. 15G and H). Comparable to the leg, the wing may be an organ that can receive signals from females. Recent research found that pheromone sensing *ppk29* and *ppk23* were significantly expressed in the wing [65], thus indicating that the wing is also an intriguing organ for pheromone sensing function and may contribute to the mating behavior of males. Future research will investigate the potential role of the wings in SMD behavior.

**Fig. 15.**
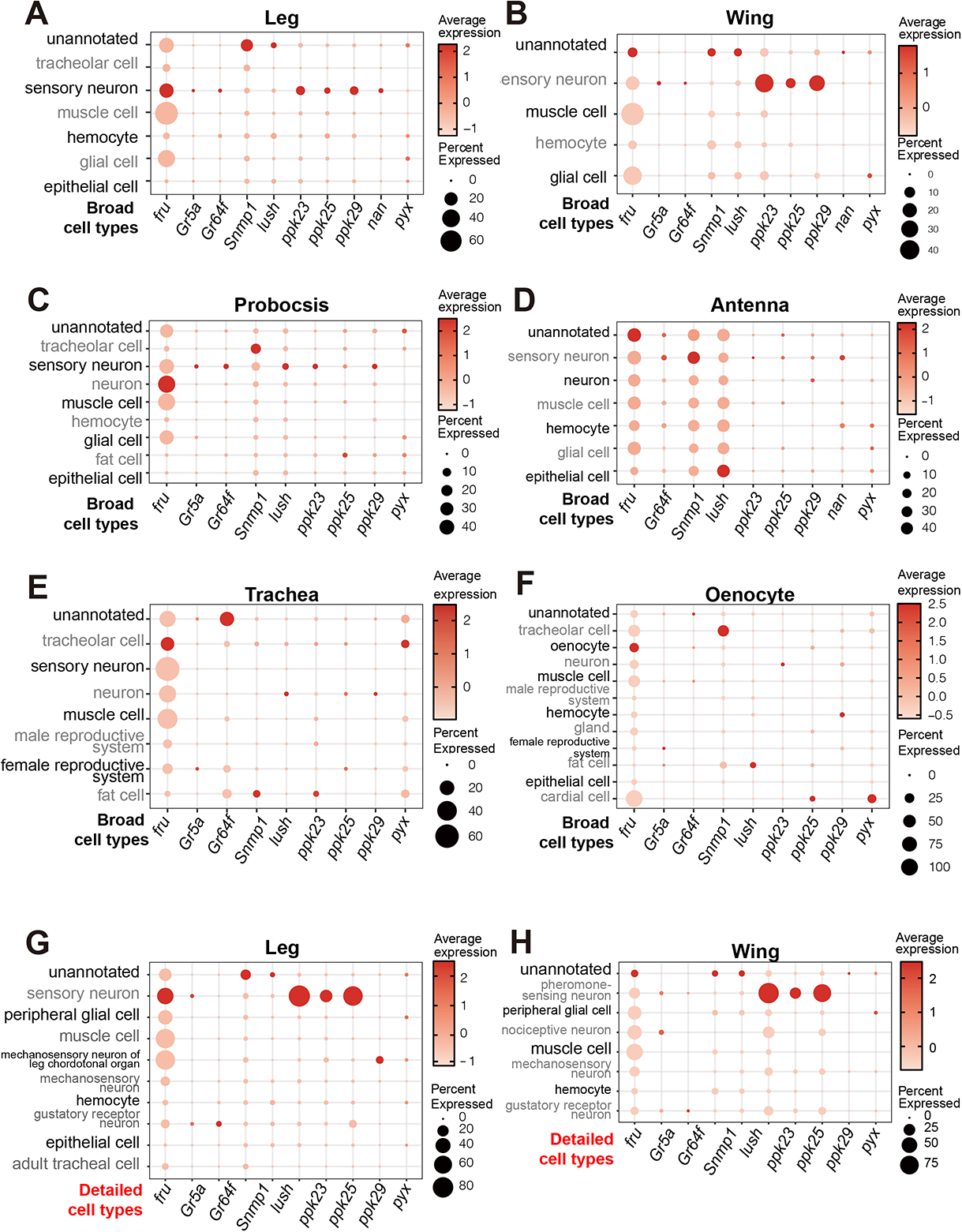
Dot plot of the 10 genes involved in SMD in each cell type in different tissues (A-H) The size of dots represents the percentagew of cells in one cell type expressing the gene of interest; the intensity of color reflects the average scaled expression. We used broad annotation for (A-F) and detailed one for (G) and (H).

*Pyrexia* has been identified as a new thermal transient receptor potential channel that endows tolerance to high temperatures in *D. melanogaster* [66] although its function in the leg is not well characterized [67]. For appropriate proprioception and hearing, *Drosophila* recruits a special sensory system, the chordotonal organ (ChO) [50, 68]. The expression of PYX in the ChO protects flies from rapid paralysis at 40°C [66]. PYX also performs an important temperature-independent function known as geotactic behavior [69, 70]. Interestingly, *pyx* is required to synchronize clock neurons in the brain [71]; however, its function has been observed in the cap cells of Johnston’s organ in the antenna, but not the leg. Thus, we suggest that the expression of PYX in the male’s sugar receptor neurons also modulates the function of clock neurons and might function as an important pacemaker for SMD behavior.

Our screening of receptors involved in proprioception, hearing, mechanosensation, and temperature sensation identified that only *nanchung* and *pyrexia* in *Gr5a*-positive sugar cells were required to provide appropriate sensory modality for SMD behavior (Fig. 10, Fig. S10, Fig. 11, and Fig. S11). We also showed that the auditory/proprioception mutant *iav^1^* did not exhibit SMD behavior (Fig. 2F). *Nanchung* and *inactive* are two independent TRPV channel subunits and mediate hearing in *Drosophila* [72]. *Nanchung* is also required for hygrosensation [73]. Recently, Lone at al. reported that mechanosensory stimulation *via nanchung*-expressing neurons can induce daytime sleep [74], thus suggesting that *nanchung* mediates orbital motion-induced sleep which is closely associated to the circadian clock. Although we cannot rule out the possible involvement of other mechanosensory receptors on SMD behavior, deciphering the neural circuits that mediate mechanosensation and mating duration will be an important aspect of future research.

In summary, we report a novel sensory pathway that controls mating investment related to sexual experiences in *Drosophila*. Since both LMD and SMD behaviors are involved in controlling male investment by varying the interval of mating, these two behavioral paradigms will provide a new avenue to study how the brain computes the ‘interval timing’ that allows an animal to subjectively experience the passage of physical time [75–80].

## AUTHOR CONTRIBUTION

SGL performed MD assays, data analysis, and mapping leg cells by live imaging analysis as shown in Fig. 3, Fig. 4, Fig. S4, Fig. 5, Fig. S5, Fig. 6, Fig. S6, Fig. 7I, Fig. 8, Fig. S8, Fig. 9, Fig. S9, Fig. 11 and Fig. S12. CK and TNS performed the evolutionary modeling shown in Fig. 13 and Box. S1. BS, KHN, AGP and DB performed the MD assays and data analysis shown in Fig. 10 and Fig. S10. AHA and BT performed male reproductive fitness experiments and the data analysis shown in Fig. 13. AM and MZ performed MD assays and the data analysis shown in Fig. 11. ACA, MEI, KW and TS performed MD assays and data analysis for the control experiments shown in Fig. S13 and Fig. S14. EL performed male leg imaging and the data analysis shown in Fig. S11 and designed the cover image for this manuscript. DS performed the MD assays and data analysis related to the temporal activation or inhibition of Gr5a neurons shown in Fig. 12. HM performed CalexA experiments and the quantification of GFP signals is shown in Fig. 12. ZW analyzed the fly cell atlas scRNA sequencing data shown in Fig. 15. WJK designed the experiments and performed MD assays, data analysis, imaging analysis, and the fly SCope data analysis shown in Fig. 1, Fig. S1, Fig. 2, Fig. S2, Fig. 3, Fig. S3, Fig. 4, Fig. S4, Fig. 5, Fig. S5, Fig. 7, Fig. S7, Fig. S11, Fig. S12, Fig. 13, Fig. 14, Fig. S15, Fig. S16 and Fig. S17, and wrote and revised the manuscript.

## ACKNOWLEDGEMENTS

We thank Dr. Joshua Bagley (University of California, San Francisco), Dr. Kyeongjin Kang (Sungkyungkwan University, Korea), Seokjun Moon (Yonsei University), Youngjoon Kim (GIST) and Ashley Kim (Mountain View) for helpful comments and support on this paper. We are also very appreciative to the colleagues who supplied us with several fly strains; Drs. Joel D. Levine and Joshua J. Krupp (University of Toronto, Canada), Dr. Barry Dickson (HHMI Janelia Research Campus, USA), Dr. Toshiro Aigaki (Tokyo Metropoitan University, Japan), Dr. Martin Heisenberg (Universität Würzburg, Germany), Dr. Michael Gordon (University of British Columbia, Canada), Dr. Richard Benton (Université de Lausanne, Switzerland), and Dr. Ralf Stanewsky (University of Münster, Germany). This research was supported a University of Ottawa Startup grant to WJK, a University of Ottawa Brain and Mind Research Institute/Center for Neural Dynamics Open call project grant to WJK, a University of Ottawa Interdisciplinary Research Group Funding Opportunity (IRGFO stream 1 and 2) Grant to WJK, a Natural Sciences and Engineering Research Council of Canada (NSERC) Discovery grant (reference: 211406) to WJK, a Mitacs Globalink Research Internship Program grant to WJK, and Startup funds from HIT Center for Life Science to WJK. This research was also supported by the Brain Pool Program of the National Research Foundation in Korea to WJK, Burroughs Wellcome Fund Collaborative Research Travel Grants (reference: 1017486) to WJK and a NVIDIA Academic Hardware Grant Program to WJK. The authors would like to express their gratitude to EditSprings (https://www.editsprings.cn) for the expert linguistic services provided.

## EXPERIMENTAL PROCEDURES

### Fly Rearing and Strains

*Drosophila melanogaster* were raised on cornmeal-yeast medium at similar densities to yield adults with similar body sizes. Flies were kept in 12 h light: 12 h dark cycles (LD) at 25°C (ZT 0 is the beginning of the light phase, ZT12 beginning of the dark phase) except for some experimental manipulation (experiments with the flies carrying tub-GAL80ts). Wild-type flies were Canton-S. To reduce the variation from genetic background, all flies were backcrossed for at least 3 generations to CS strain. All mutants and transgenic lines used here have been described previously.

We are very grateful to the colleagues who provided us with many of the lines used in this study. We obtained the following lines from Dr. Joel D. Levine and Joshua J. Krupp (University of Toronto, Canada): *PromE(800)-GAL4* (*oeno-GAL4* in this study); from Dr. Barry Dickson (HHMI Janelia Research Campus, USA): *UAS[stop]mCD8GFP; fruFLP*, *UAS[stop]nsybGFP; fruFLP, UAS[stop]TNTactive; fruFLP*, *fru-GAL4*; from Dr. Toshiro Aigaki (Tokyo Metropoitan University, Japan): *UAS-mSP*; from Dr. Martin Heisenberg (Universität Würzburg, Germany): *WT Berlin, ninaE17*; from Dr. Michael Gordon (University of British Columbia, Canada): *Gr5a-lexA, Gr64f-lexA, ppk23-GAL4, ppk25-GAL4, ppk29-GAL4*; from Dr. Richard Benton (Université de Lausanne, Switzerland): *Ir84a^-/-^*; from Dr. Ralf Stanewsky (University of Münster, Germany): *pyx-GAL4*.

The following lines were obtained from Bloomington Stock Center (#stock number): *Orco^1^* (#23129), *Orco^2^* (#23130), *UAS-tubGAL80^ts^* (#7018), *Df(1)^Exel6234^* (#7708), *UAS-tra^F^* (#4590), *GustD^x6^* (#8607), *Gr66a-GAL4* (#28801), *UAS-mCD8GFP* (#5130), *UAS-RedStinger* (#8547), *snmp1^1^* (#25043), *snmp1^2^* (#25042), *UAS-snmp1* (#25044), *Snmp2-RNAi* (#51432), *UAS-TNT* (#28997), *UAS-dicer* (#24650, #24651), *tra-RNAi* (#28512), *lush-RNAi* (#31657), *tud^1^* (*son-of-tudor* males were the sons of Oregon R males and virgin tudor females: tud1 bw sp/tud1 bw sp) (#1786), *lexAop-tdTomato.nls*, *UAS-Stinger* (#66680), *Gr5a-RNAi* (#31282), *Gr43a-RNAi* (#64881), *Gr61a-RNAi* (#54030), *Gr64c-RNAi* (#36734), *ppk23-RNAi* (#28350), *ppk25-RNAi* (#27088), *ppk29-RNAi* (#27241), *pkd2-RNAi* (#31296, #31675), *nan-RNAi* (#31295, #31674, #53312), *iav-RNAi* (#25865), *norpA-RNAi* (#31113, 31197), *nompC-RNAi* (#31512, 31689), *pyx-RNAi* (#31297, 51836), *pain-RNAi* (#31510, #31511, #61299), *wtrw-RNAi* (#31292, #31293, #51503), *trp-RNAi* (#61352, #31649, #64932), *TrpA1-RNAi* (#31384, #31504, #31384), *Trpml-RNAi* (#31673, #31294), *Trpm-RNAi* (#51731, #57871), *Trpγ-RNAi* (#31298, #53313), *TMC-RNAi* (#50984), *fru-lexA* (#66698), *Gr64f-GAL4* (#57668), *se^1^*(*sepia* mutants for fecundity test, #1668), *elav^c155^; UAS-Dcr-2* (#25750), CalexA (#66542), *UAS-TrpA1* (#26264), *UAS-shi^ts^* (#44222); from Vienna Drosophila Stock Center: *Gr5a-RNAi* (#v13730), *Gr43a-RNAi* (#v39518), *Gr61a-RNAi* (#v106007), *Gr64a-RNAi* (#v103342), *Gr64b-RNAi* (#v42517), *Gr64d-RNAi* (#v29422), *Gr64e-RNAi* (#v109176), *Gr64f-RNAi* (#v105084). Following transgenic stocks are available from Korea Drosophila Resource Center (KDRC): *UAS[stop]TNTinactive; fruFLP* (1124).

### Mating Duration Assays

Mating duration assay was performed as previously described [14, 17]. For naïve males, 4 males from the same strain were placed into a vial with food for 5 days. For experienced males, 4 males from the same strain were placed into a vial with food for 4 days then eight CS virgin females were introduced into vials for last 1 day before assay. Five CS females were collected from bottles and placed into a vial for 5 days. These females provide both sexually experienced partners and mating partners for mating duration assays. At the fifth day after eclosion, males of the appropriate strain and CS virgin females were mildly anaesthetized by CO2. After placing a single female in to the mating chamber, we inserted a transparent film then placed a single male to the other side of the film in each chamber. After allowing for 1 h of recovery in the mating chamber in a 25°C incubator, we removed the transparent film and recorded the mating activities. Only those males that succeeded to mate within 1 h were included for analyses. Initiation and completion of copulation were recorded with an accuracy of 10 sec, and total mating duration was calculated for each couple. All assays were performed from noon to 4 pm. We conducted blinded studies for every test.

### Courtship Assays

Courtship assay was performed as previously described [90], under normal light conditions in circular courtship arenas 11 mm in diameter, from noon to 4 pm. Courtship latency is the time between female introduction and the first obvious male courtship behavior such as orientation coupled with wing extensions. Once courtship began, courtship index was calculated as the fraction of time a male spent in any courtship-related activity during a 10 min period or until mating occurred. Mating initiation is the time after male flies successfully mounted on female.

### Locomotion Assays

For climbing assay, individual flies were placed in a 15 ml falcon tube (Fisher Scientific) and were gently tapped to the bottom of the tube. The time taken for the flies to climb 8 cm of the tube wall was recorded. Each fly was tested 5 times. Other than a single instance, all flies were seen to reach the target height within 2 min, which was the experimental cut-off time. Velocity was obtained by dividing the lines (mm) a fly crossed (distance walked) by time (sec) a fly reached the line of the tube. For horizontal (spontaneous) locomotor activities, a single fly was first brought to the middle of the column by gentle shaking and then the fly movement was constantly monitored for 5 min and recorded. Total fraction of time flies walked during 5 min was calculated and number of stops during 5 min was also counted then calculated [91].

### Immunostaining and Antibodies

As described before [17], brains dissected from adults 5 days after eclosion were fixed in 4% formaldehyde for 30 min at room temperature, washed with 1% PBT three times (30 min each) and blocked in 5% normal donkey serum for 30 min. The brains were then incubated with primary antibodies in 1% PBT at 4oC overnight followed with fluorophore-conjugated secondary antibodies for 1 hour at room temperature. Brains were mounted with anti-fade mounting solution (Invitrogen, catalog #S2828) on slides for imaging. Primary antibodies: chicken anti-GFP (Aves Labs, 1:1000), rabbit anti-DsRed express (Clontech, 1:250), mouse anti-Bruchpilot (nc82) (DSHB, 1:50), mouse anti-PDF (DSHB, 1:100). Fluorophore-conjugated secondary antibodies: Alexa Fluor 488-conjugated goat anti-chicken (Invitrogen, 1:100), Alexa Fluor 488-conjugated donkey anti-rabbit (Invitrogen, 1:100), RRX-conjugated donkey anti-rabbit (Jackson Lab, 1:100), RRX-conjugated donkey anti-mouse (Jackson Lab, 1:100), Dylight 649-conjugated donkey anti-mouse (Jackson Lab, 1:100).

### Quantitative Analysis of GFP Fluorescence

To quantify the calcium level in leg sensory neurons, we measured fluorescence intensity using the measure tool of ImageJ (National Institutes of Health, http://rsb.info.nih.gov/ij). Fluorescence was quantified in a manually set region of interest (ROI) of the 4^th^ or 5^th^ tarsus. To compensate for differences in fluorescence between different ROI, GFP fluorescence for CaLexA was normalized to autofluorescence, and then the fluorescence of ROI was quantified using the measure tool of ImageJ. All specimens were imaged under identical conditions.

### Statistical Analysis

Statistical analysis of mating duration assay was described previously [14, 17]. More than 36 males (naïve or experienced) were used for mating duration assay. Our experience suggests that the relative mating duration differences between naïve and experienced condition are always consistent; however, both absolute values and the magnitude of the difference in each strain can vary. So, we always include internal controls for each treatment as suggested by previous studies [92]. Therefore, statistical comparisons were made between groups that were naively reared or sexually experienced within each experiment. As mating duration of males showed normal distribution (Kolmogorov-Smirnov tests, p > 0.05), we used two-sided Student’s t tests. Each figure shows the mean ± standard error (s.e.m) (*** = p < 0.001, ** = p < 0.01, * = p < 0.05). All analysis was done in GraphPad (Prism). Individual tests and significance are detailed in figure legends.

When we compare the difference of mating duration in experiments without internal control built in, we always performed control experiments of wild type for each independent experiment for internal comparison. And in this case, we analyzed data using ANOVA for statistically significant differences (at a 95.0% confidence interval) between the means of mating duration for all conditions. If a significant difference between the means was found by Kruskal-Wallis test, then the Dunn’s Multiple Comparison Test was used to compare the mean mating duration of each condition to determine which conditions were significantly different from condition of interest. (# = p < 0.05)

Besides traditional t-test for statistical analysis, we added estimation statistics for all MD assays and two group comparing graphs. In short, ‘estimation statistics’ is a simple framework that— while avoiding the pitfalls of significance testing—uses familiar statistical concepts: means, mean differences, and error bars. More importantly, it focuses on the effect size of one’s experiment/intervention, as opposed to significance testing [93]. In comparison to typical NHST plots, estimation graphics have the following five significant advantages such as (1) avoid false dichotomy, (2) display all observed values (3) visualize estimate precision (4) show mean difference distribution. And most importantly (5) by focusing attention on an **effect size**, the difference diagram encourages quantitative reasoning about the system under study [94]. Thus, we conducted a reanalysis of all of our two group data sets using both standard t-tests and estimate statistics. In 2019, the Society for Neuroscience journal eNeuro instituted a policy recommending the use of estimation graphics as the preferred method for data presentation [95].

### Egg and Progeny Counting

We performed egg laying assay as previously described [18]. After we count the number of eggs, we kept vials in 25°C incubator and counted the total number of progenies ecolsed from them.

### Fecundity Test by Introducing the Second Male

Basically, we followed the protocols previously described by other group [96]. In short, *se^1^* or CS virgin females were introduced to *se^1^* or CS males either as naïve or experienced condition for 24 hours to be confident of all females’ mating with the first males. Then we introduced the second males for 24 hours. After this treatment, we separated females from second males then counted the number of progenies from females. To confirm that the effect from this fecundity test was not originated from the genotype background, we performed the same experiments by reversing the genotypes of the first and second males (*se^1^* then CS vs. CS vs. *se^1^*). We calculated the percentage of progeny either from the first male or the second male by counting the eye color of progeny.

### Single-nucleus RNA-sequencing Analyses - Data and Code Availability

snRNAseq dataset analyzed in this paper is published in [64] and available at the Nextflow pipelines (VSN, https://github.com/vib-singlecell-nf), the availability of raw and processed datasets for users to explore, and the development of a crowd-annotation platform with voting, comments, and references through SCope (https://flycellatlas.org/ scope), linked to an online analysis platform in ASAP (https://asap.epfl.ch/fca).

### Gene expression pattern analyses in different tissues

For the gene expression pattern of the 10 genes involved in SMD in each cell type of leg and other tissues, we used the single-cell RNA-seq data from https://flycellatlas.org [64], and the 10x Genomics stringent loom files were downloaded. The cell types are split by FCA.

The digital expression matrices were analyzed with the Seurat 4.1.0 R package [97]. The dot plots of the 10 genes involved in SMD in each cell type of different tissues were then made using the ‘DotPlot’ function with broad annotation (broad cell types) and the annotation (detailed cell types), respectively.

## SUPPLEMENTARY FIGURE LEGENDS

**Fig. S1.**
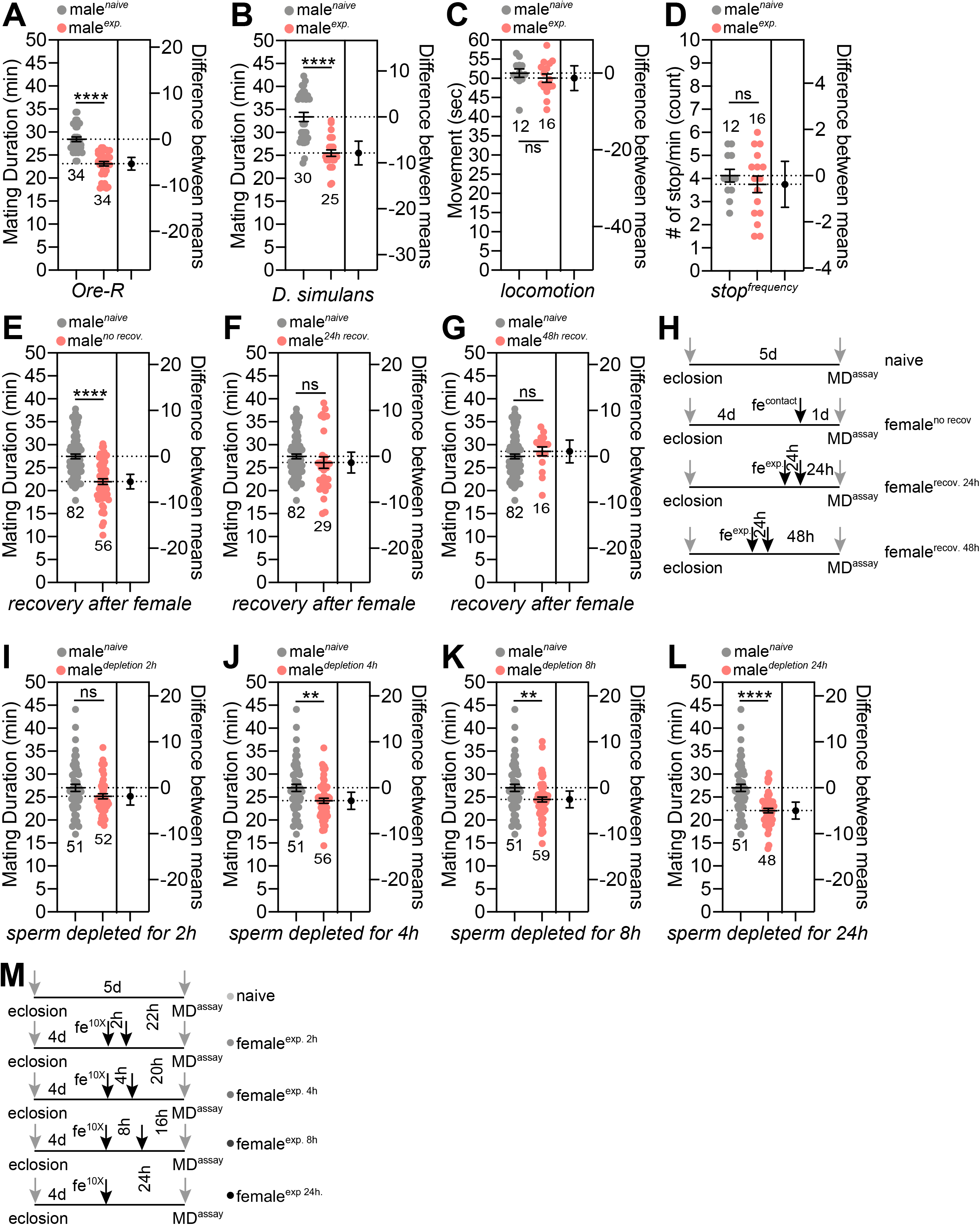
General characteristics of ‘Shorter-Mating-Duration (SMD)’ behaviour. (A) Mating duration (MD) assays of Oregon-R males and (B) *Drosophila simulans* males (C) Locomotion of naïve and experienced male flies were quantified as velocity by locomotion activity by horizontal paradigm, and (D) stop frequency by horizontal paradigm. See **EXPERIMENTAL PROCEDURES** section for detailed methods. (E-G) MD assays of CS males after isolated from female experience. Males were reared with sufficient numbers of virgin females for 24 h to be assured having sexual experience then isolated. Assay times after isolation are below the boxes as (E) no recovery, (F) 24 h recovery, and (G) 48 recovery. (H) The diagram of MD assays of CS males after different time of isolation after sexual experience with females. (I-L) MD assays of CS males after sperm deleted as shown in (M).

**Fig. S2.**
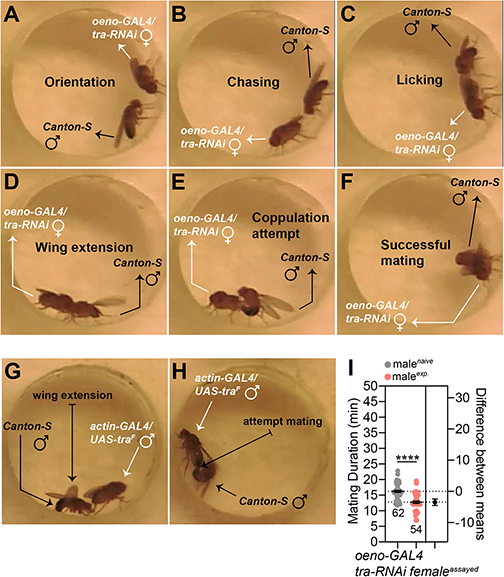
Sensory inputs required for inducing SMD behavior. (A) CS male court oenocytes-musicalized female and show orientation behavior, (B) chasing (C) licking (D) wing extension, (E) copulation attempt, and (F) can successfully mate with it. (G) CS male court feminized male and show wing extension behavior and (H) copulation attempt. (I) MD assays of CS males with oenocytes-masculinized female as a female partner to test whether genotypes of female partners affect MD.

**Fig. S3.**
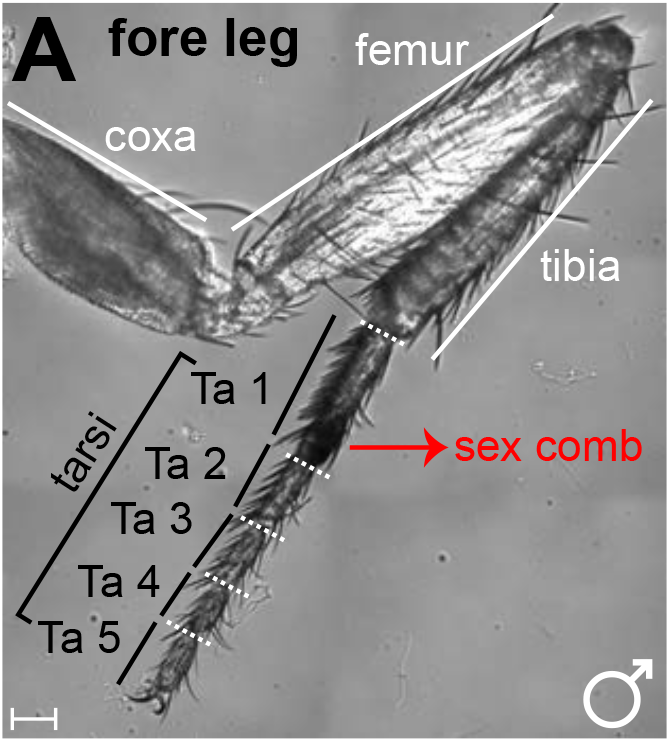
The foreleg of male *Drosophila melanogaster* (A) The anatomical structures of male foreleg are labeled. Ta1-Ta5 comprise fore tarsus and represents tarsomeres 1-5, respectively.

**Fig. S4.**
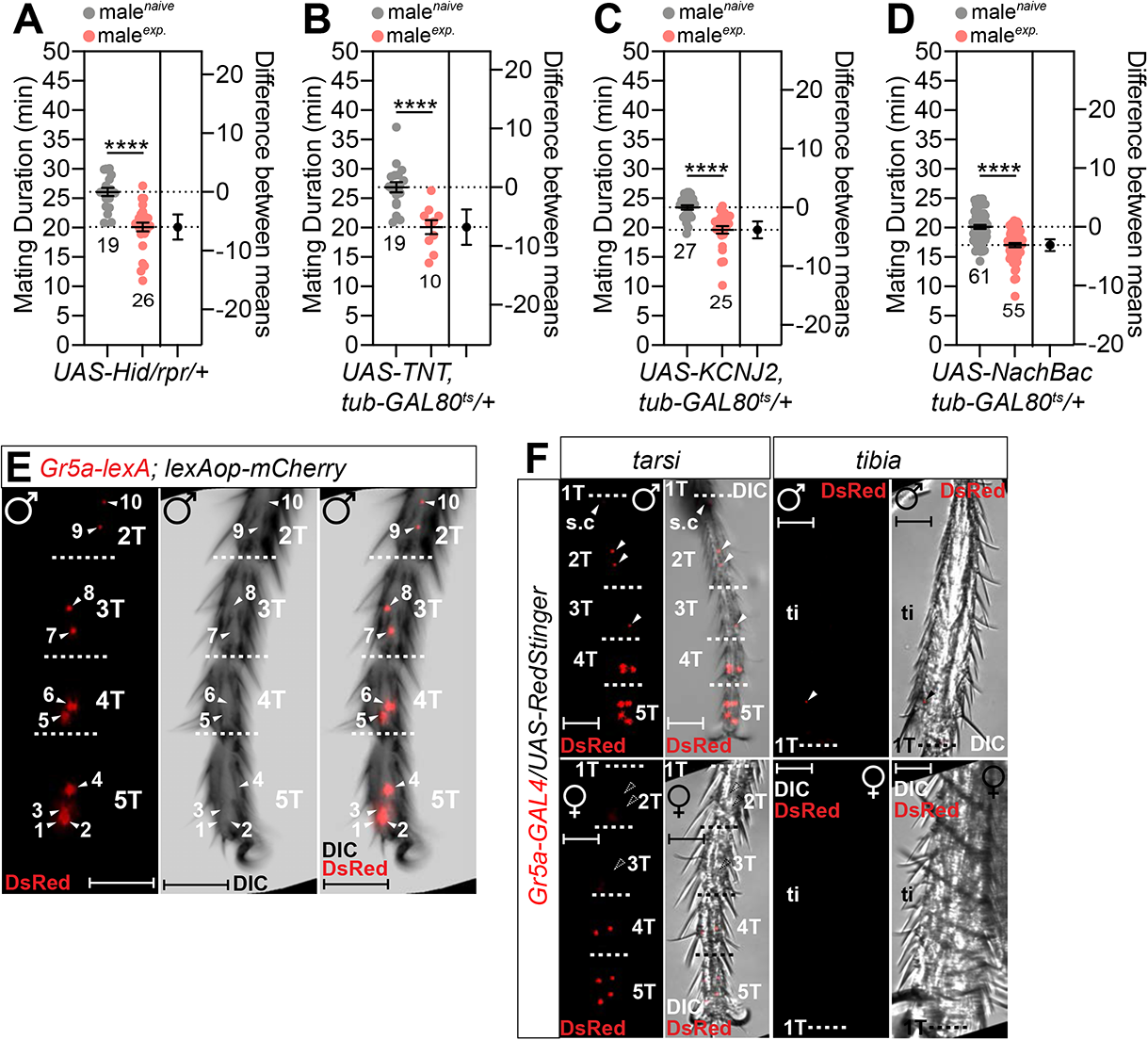
Control experiments for MD assays in Fig. 4 and the location of *Gr5a*-positive neurons in male foreleg (A-D) MD assays of (A) *UAS-Hid/rpr* (B) *UAS-TNT, tub-GAL80^ts^* (C)*UAS-KCNJ2, tub-GAL80^ts^* (D) *UAS-NachBac, tub-GAL80^ts^* crossed with CS. (E) Foreleg tarsus of male flies expressing *Gr5a-lexA* together with *lexAOP-mCherry*. White arrows indicate *Gr5a-*positive neurons and numbers represent the order from the distal part of the leg. (F) Foreleg tarsus (left panels) and tibia (right panels) of male (top panels) or female (bottom panels) flies expressing *Gr5a-GAL4* together with *UAS-RedStigner*. White arrows indicate *Gr5a-*positive neurons. White arrows with dotted line indicate missing neurons in female leg compared to male leg.

**Fig. S5.**
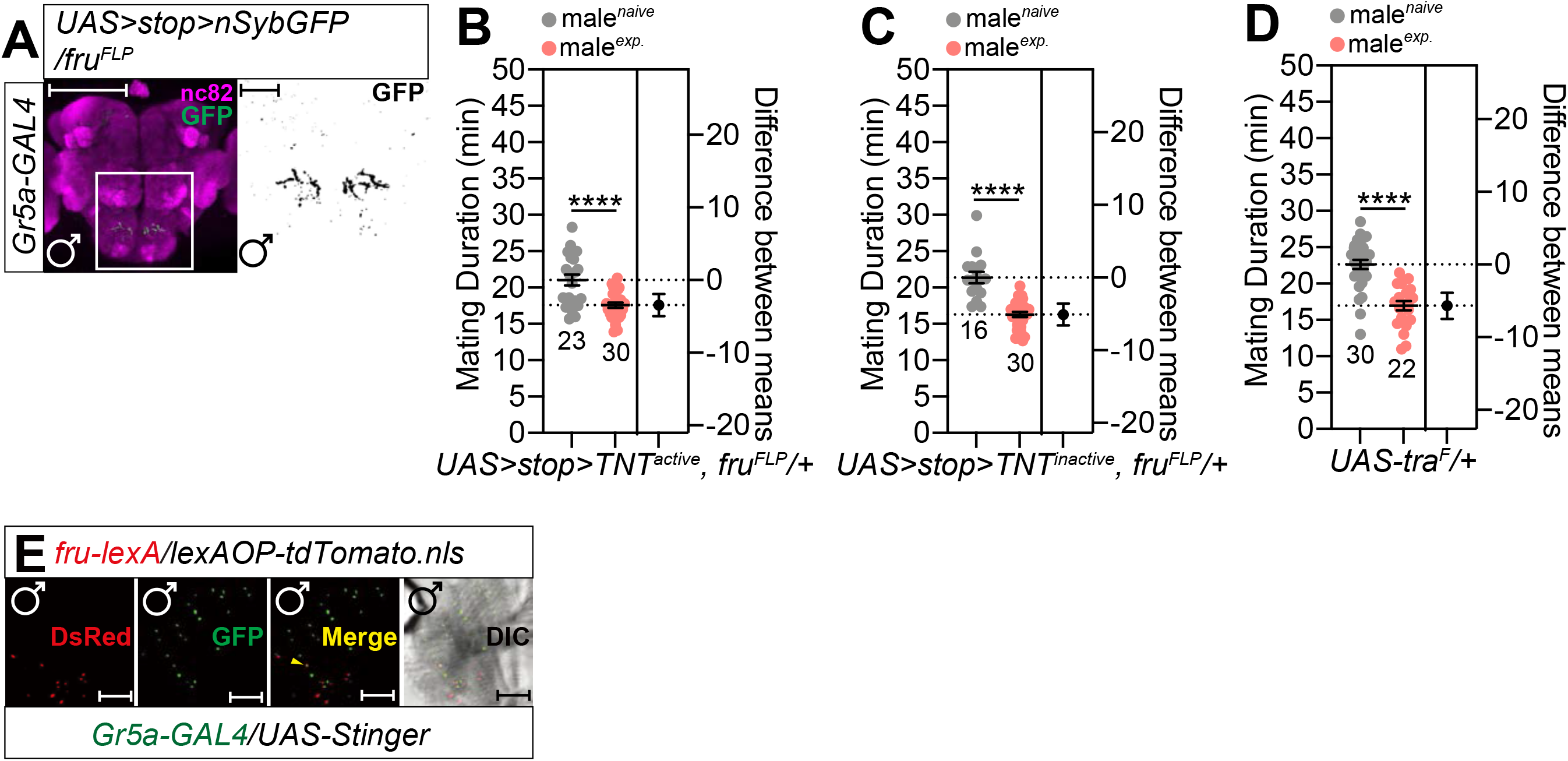
(A) Brains of male flies expressing *Gr5a-GAL4* together with *UAS>stop>nSybGFP; fru^FLP^* were immunostained with anti-GFP (green) and nc82 (magenta) antibodies. Scale bars represent 100 μm in the colored panels and 10 μm in the grey panels. White boxes indicate the magnified regions of interest presented next right panels. The right panels are presented as grey scale for clearly showing the axon projection patterns of gustatory neurons in the adult subesophageal ganglion (SOG) labeled by *GAL4* drivers. (B-D) Control experiments for MD assays in Fig. 5. MD assays of (B) *UAS>stop>TNT_active_; fru^FLP^* (C) *UAS>stop>TNT_inactive_; fru^FLP^* (D) *UAS-tra^F^* crossed with CS. (E) Proboscis of male flies expressing *fru-lexA; lexAOP-tdTomato.nls*, *UAS-Stinger* with *Gr5a-GAL4* were imaged in live. Scale bars represent 100 μm.

**Fig. S6.**
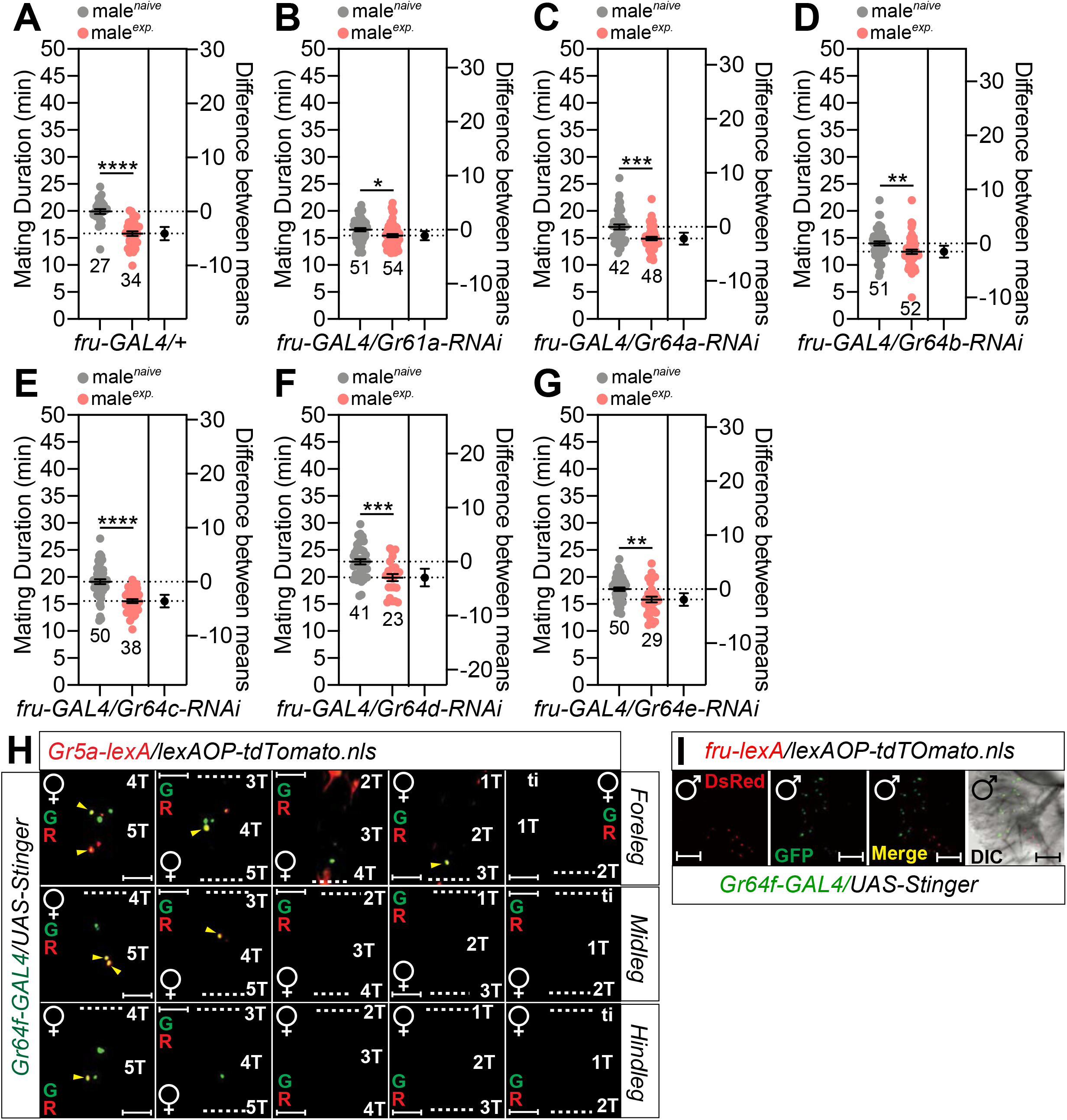
(A) Control experiments for MD assays in Fig. 6B-D and **Fig. S6B-G**. (B-G) MD assays of flies expressing *fru-GAL4* driver together with (B) *Gr61a-RNAi* (C) *Gr64a-RNAi* (D) *Gr64b-RNAi* (E) *Gr64c-RNAi* (F) *Gr64d-RNAi* (G) *Gr64e-RNAi.* (H) Female foreleg (upper panels), midleg (middle panels), and hindleg (bottom panels) of flies expressing *Gr5a-lexA* and *Gr64f-GAL4* drivers together with *lexAOP-tdTomato* and *UAS-Stinger* were imaged live under the fluorescent microscope. Yellow arrows indicate *Gr5a-*positive and *Gr64f*-positive neurons. Scale bars represent 50 mm. (I) Male proboscis of flies expressing *fru-lexA* and *Gr64f-GAL4* drivers together with *lexAOP-tdTomato* and *UAS-Stinger* were imaged live under the fluorescent microscope. Tested gustatory sugar receptors were selected based on previous study [25].

**Fig. S7.**
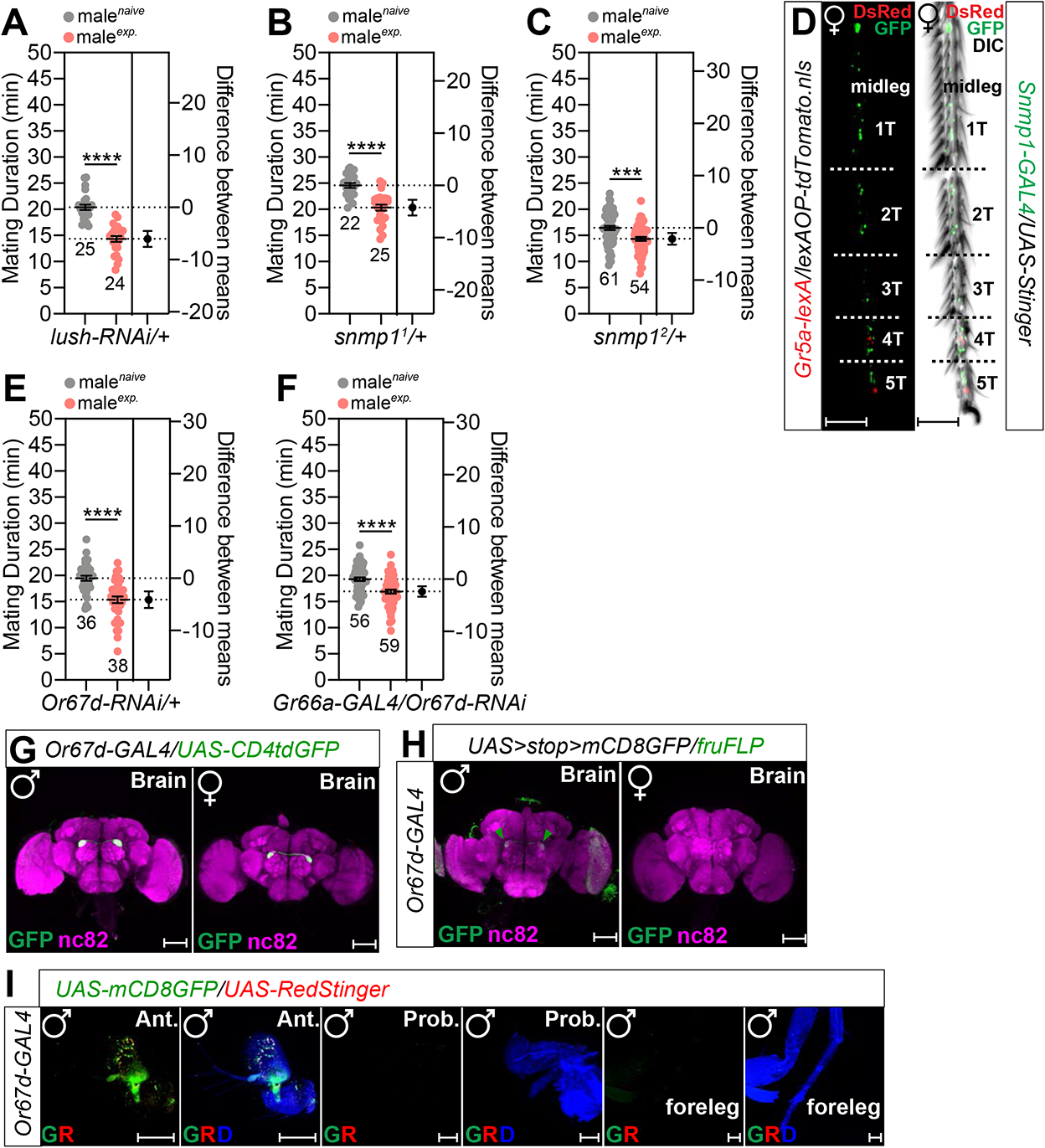
(A) Control experiments for MD assays in Fig. 7A-B. (B-C) Control experiments for MD assays in Fig. 7C-H. (D) Female foreleg of flies expressing *Gr5a-lexA* and *Snmp1-GAL4* drivers together with *lexAOP-tdTomato* and *UAS-Stinger* were imaged live under the fluorescent microscope. Scale bars represent 50 μm. (E) Control experiments for MD assays in Fig. 7K. (F) MD assays for *GAL4* mediated knockdown of *Or67d* via *Or67d-RNAi* using *Gr66a-GAL4.* (G) Brains of male (left panel) and female (right panel) flies expressing *Or67d-GAL4* together with *UAS-CD4tdGFP* were immunostained with anti-GFP (green) and nc82 (magenta) antibodies. Scale bars represent 100 μm. (H) Brains of male (left) and female (right) flies expressing *Or67d-GAL4* together with *UAS>stop>mCD8GFP; fru^FLP^* were immunostained with anti-GFP (green) and nc82 (magenta) antibodies. Scale bars represent 100 μm. (I) Male antenna (first and second panels), proboscis (third and fourth panels), and foreleg of flies expressing *Or67d-Gal4* together with *UAS-mCD8GFP, UAS-RedStinger*.

**Fig. S8.**
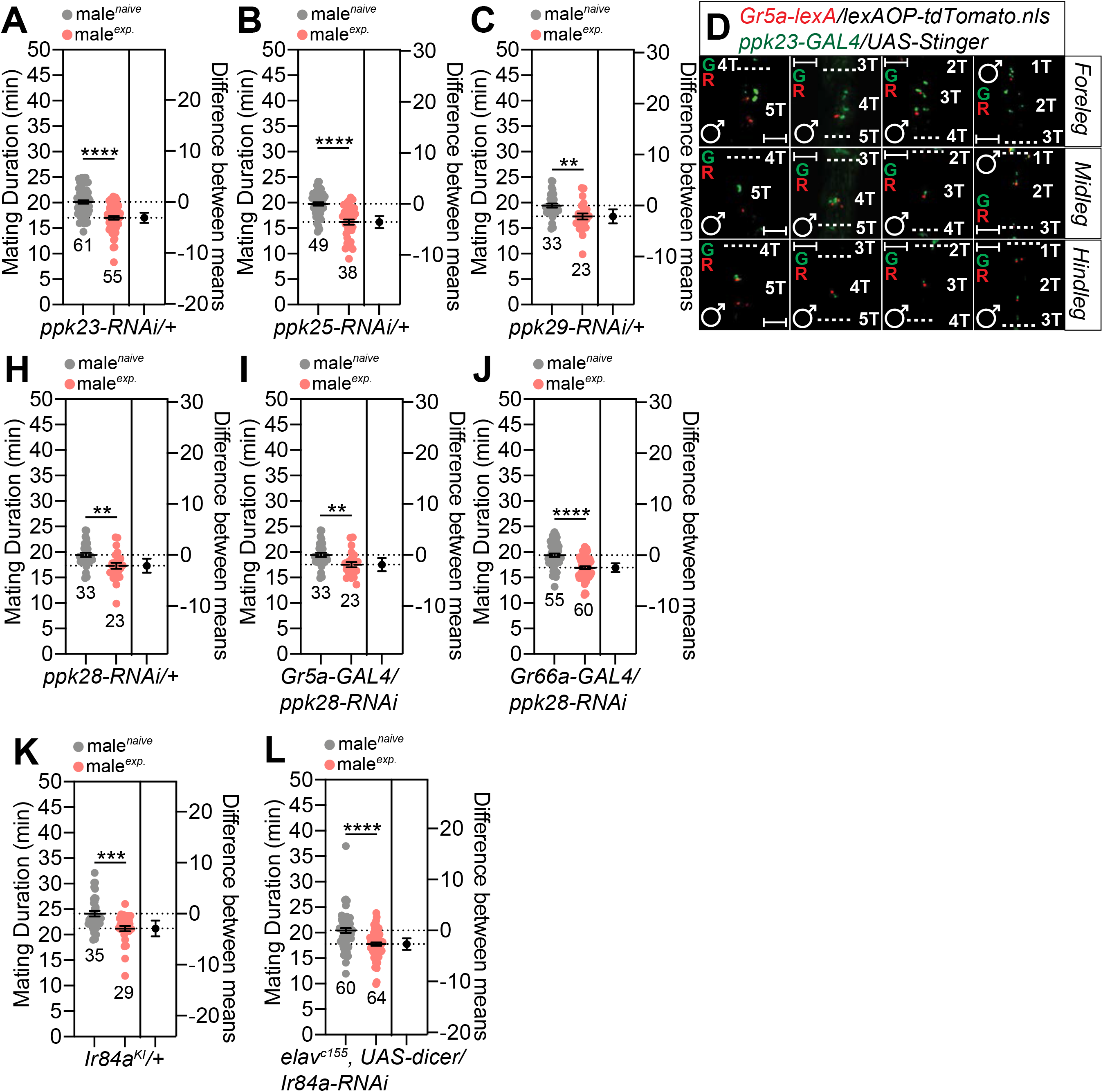
(A-C) Control experiments for MD assays in Fig. 8A-F. (D) Male foreleg (upper panels), midleg (middle panels), and hindleg (bottom panels) of flies expressing *Gr5a-lexA* and *ppk23-GAL4* drivers together with *lexAOP-tdTomato* and *UAS-Stinger* were imaged live under the fluorescent microscope. Scale bars represent 50 μm. (H) Control experiments for MD assays in **Fig. S8I-J**. (I-J) MD assays for *GAL4* mediated knockdown of PPK28 via *ppk28-RNAi* using (I) *Gr5a-GAL4* and (J) *Gr66a-GAL4* drivers. (K) Control experiments for MD assays in Fig. 8I. (L) MD assays for *GAL4* mediated knockdown of IR84a via *Ir84a-RNAi* using *elav^c155^* driver.

**Fig. S9.**
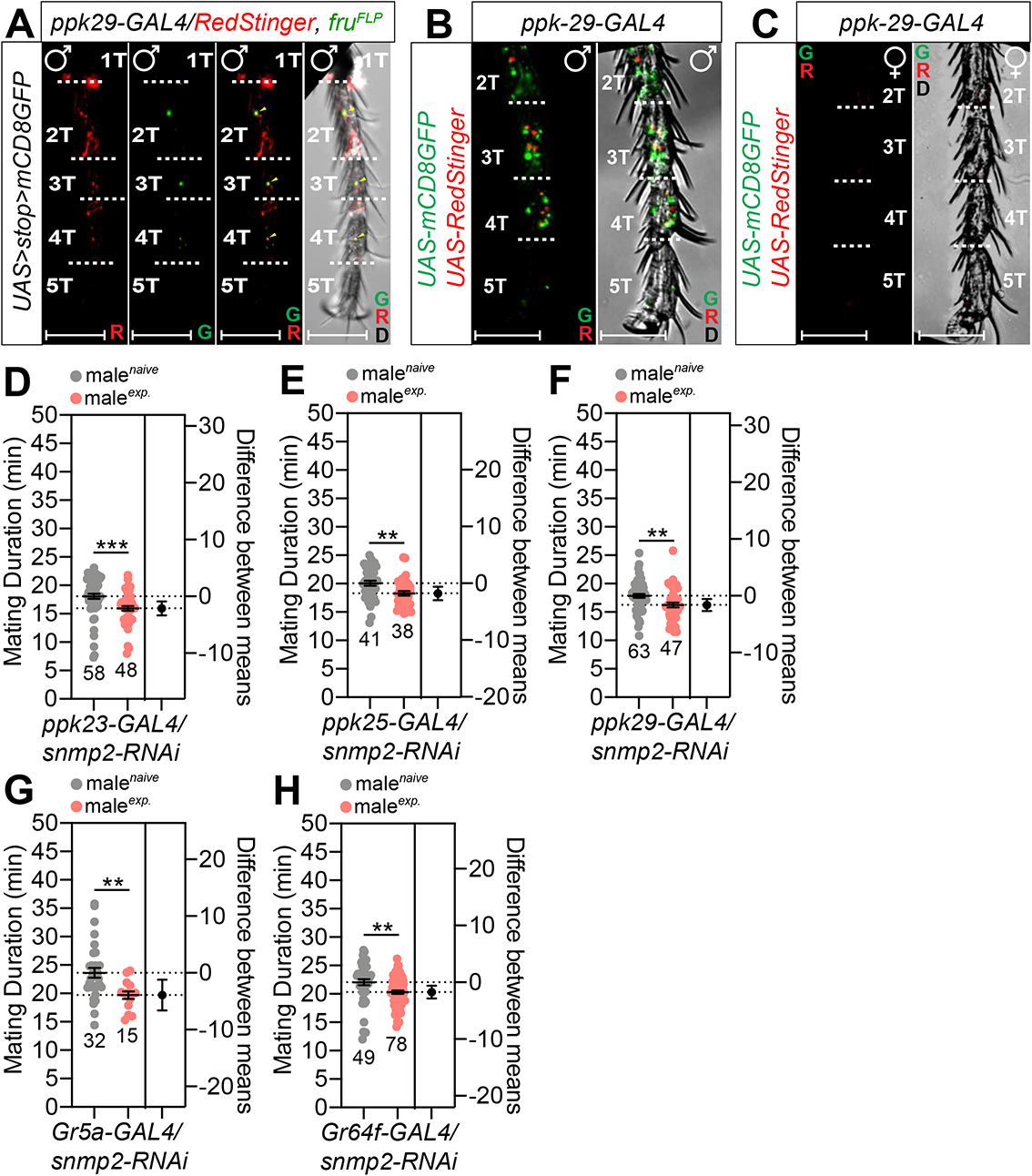
(A) Male foreleg of flies expressing *ppk29-GAL4* together with *UAS-RedStinger; UAS>stop>mCD8GFP; fru^FLP^* were imaged live under the fluorescent microscope. Scale bars represent 50 μm. (B) Male foreleg of flies expressing *ppk29-GAL4* together with *UAS-RedStinger, UAS-mCD8GFP* were imaged live under the fluorescent microscope. Scale bars represent 50 μm. (C) Female foreleg of flies expressing *ppk29-GAL4* together with *UAS-RedStinger, UAS-mCD8GFP* were imaged live under the fluorescent microscope. Scale bars represent 50 μm. (D-H) MD assays for *GAL4* mediated knockdown of SNMP2 via *snmp2-RNAi* using (D) *ppk23-GAL4*, (E) *ppk25-GAL4*, (F) *ppk29-GAL4*, (G) *Gr5a-GAL4*, and (H) *Gr64f-GAL4* drivers.

**Fig. S10.**
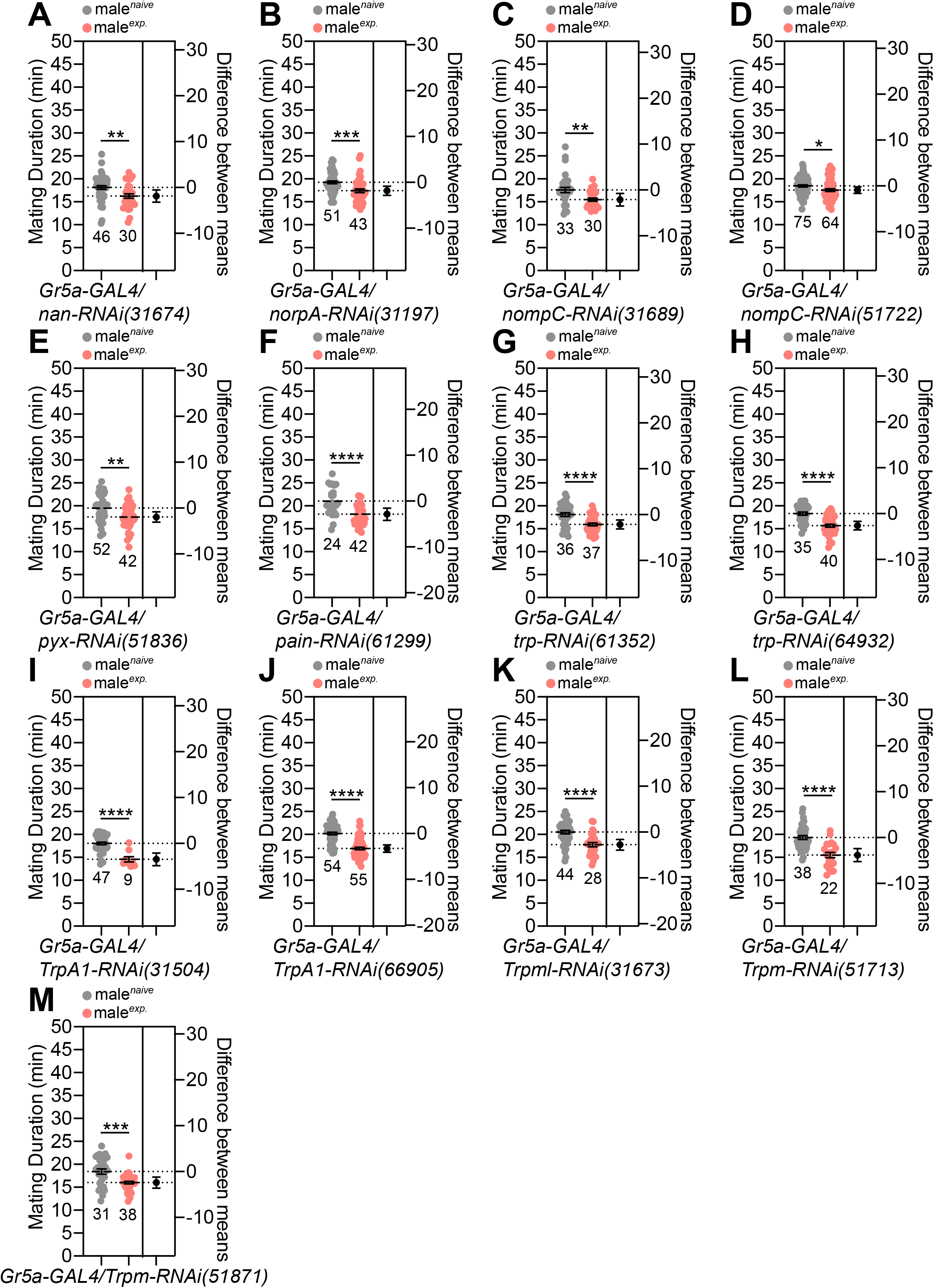
(A-M) MD assays for *GAL4* mediated knockdown of channels and receptors mediate auditory, mechanosensory, and thermal sensing using *Gr5a-GAL4* driver. Tested mechanosensory receptors were selected based on previous studies [49,51,52].

**Fig. S11.**
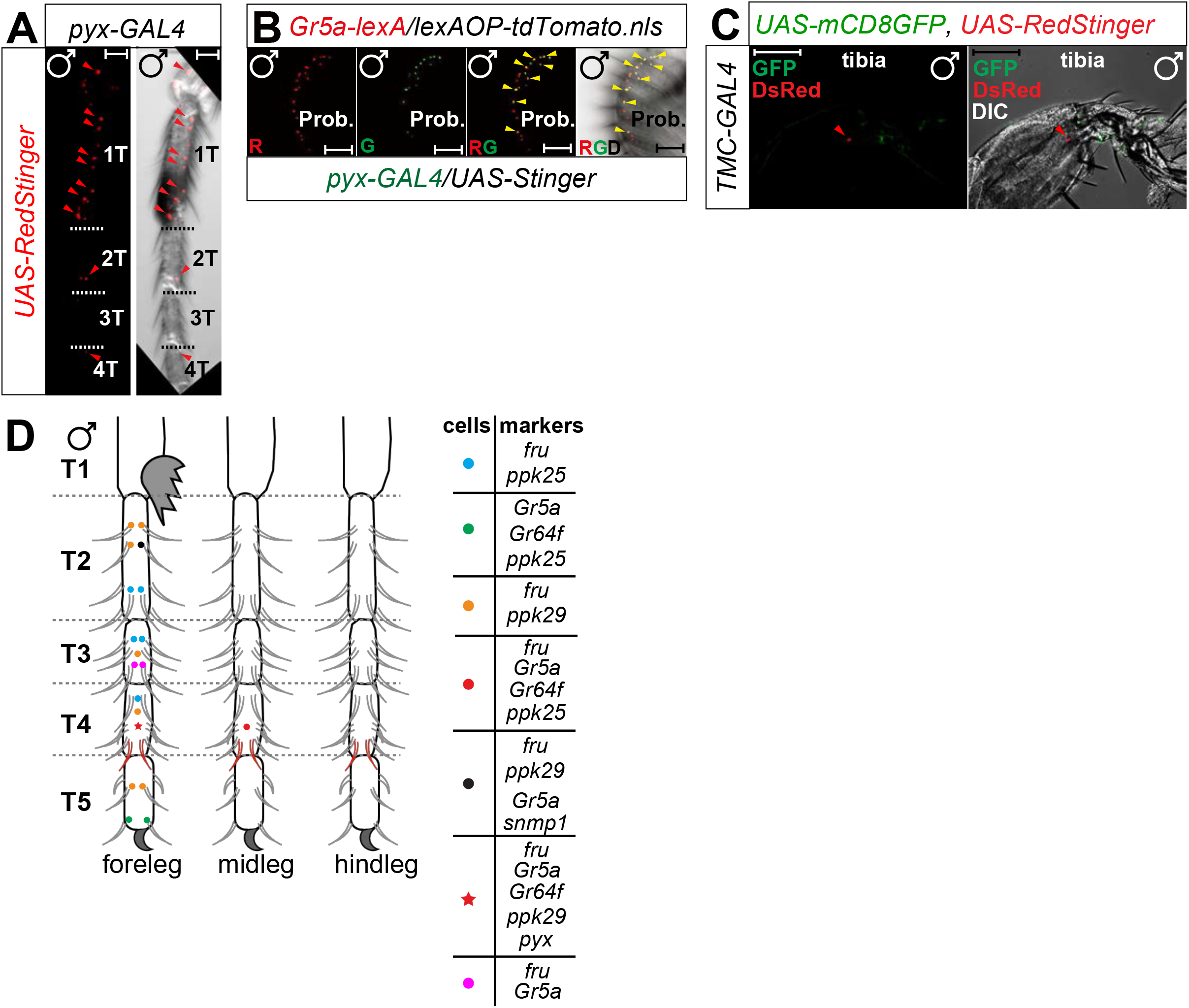
(A) Male foreleg of flies expressing *pyx-GAL4* together with *UAS-RedStinger* were imaged live under the fluorescent microscope. Red arrows indicate *pyx*-positive cells in the leg. Scale bars represent 50 μm. (B) Male proboscis of flies expressing *Gr5a-lexA* and *pyx-GAL4* drivers together with *lexAOP-tdTomato* and *UAS-Stinger* were imaged live under the fluorescent microscope. Yellow arrows indicate *Gr5a-*positive and *pyx*-positive neurons. Scale bars represent 50 μm. (C) Male foreleg tibia of flies expressing *TMC-GAL4* together with *UAS-RedStinger, UAS-mCD8GFP* were imaged live under the fluorescent microscope. Red arrow indicates the *TMC*-positive cells in tibia. Scale bars represent 50 mm. (D) A diagram of the cells in the male legs expressing genes involved in SMD behavior.

**Fig. S12.**
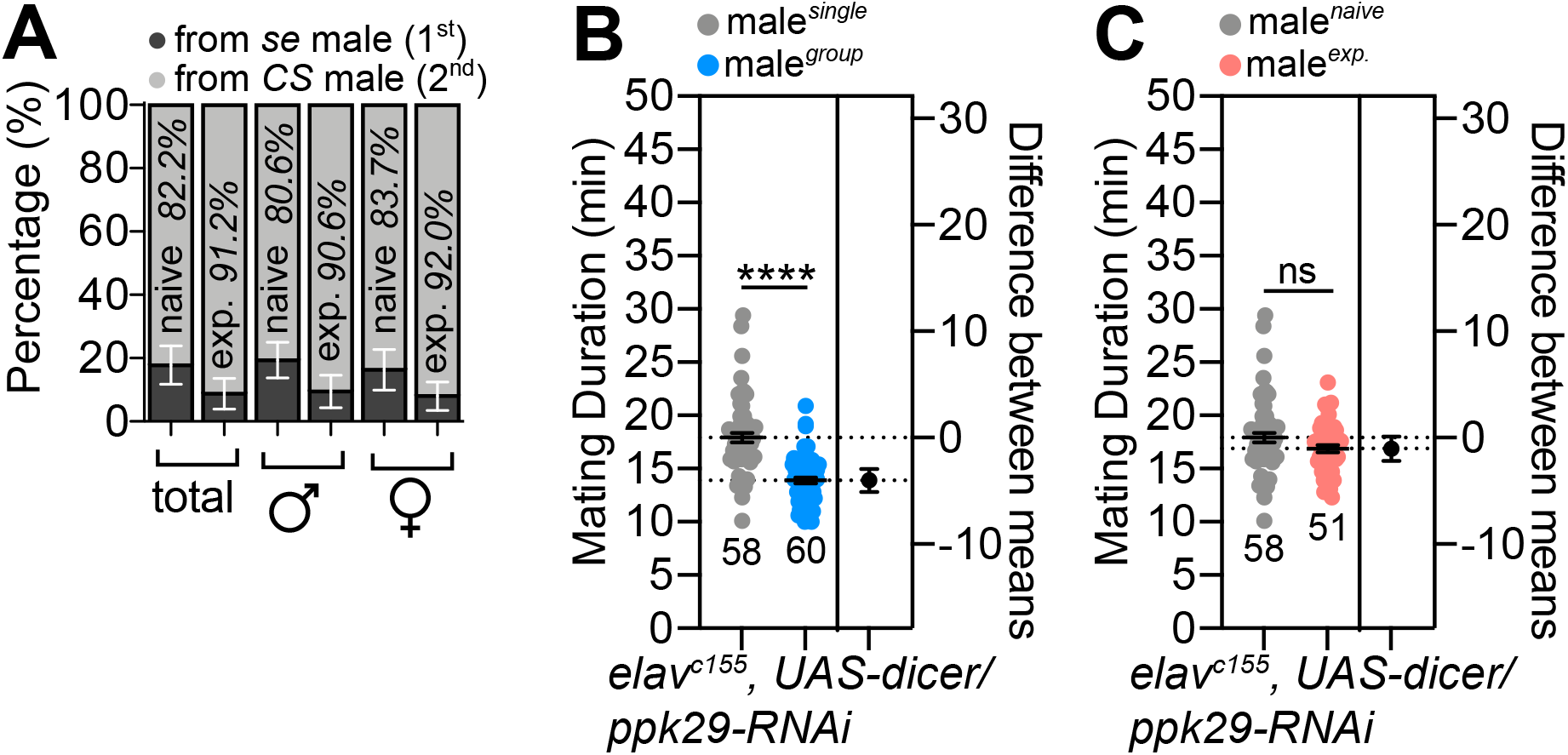
**(A)** Percentage of progeny originated from *sepia* (*se*) male vs. *CS* male. *CS* male was introduced to *se* female as first mate then followed *se* males as second mate. The eye color of progeny was counted and interpreted as the source of farther; for detailed methods, see **EXPERIMENTAL PROCEDURES**. (B-C) MD assays for *GAL4* mediated knockdown of PPK29 via *ppk29-RNAi* using *elav^c155^* for (B) LMD and (C) SMD behavior.

**Fig. S13.**
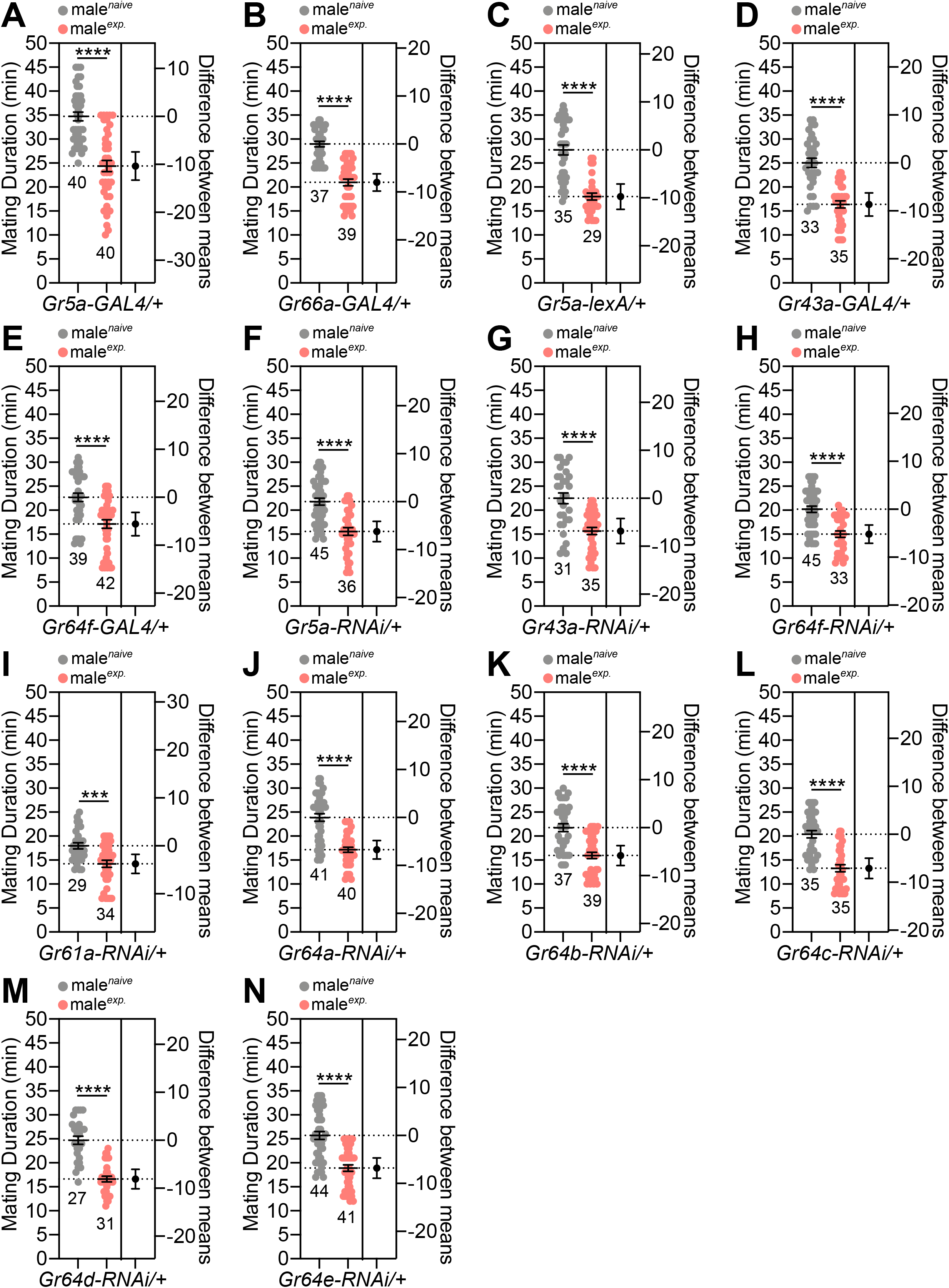
(A-N) MD assay for *GAL4, lexA*, and *RNAi* control experiments. Genotypes are labelled below the graph.

**Fig. S14.**
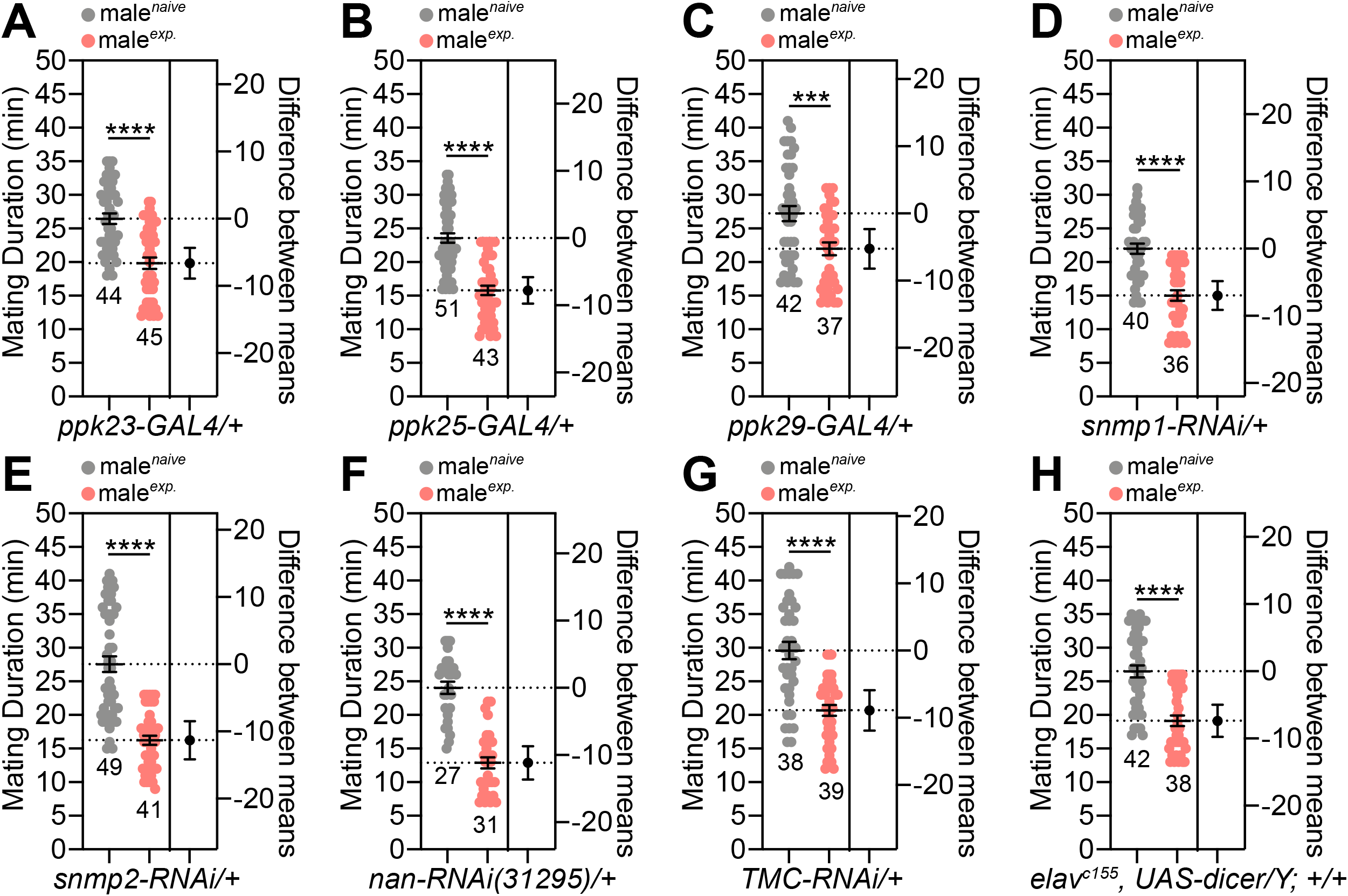
(A-H) MD assay for *GAL4, RNAi*, and *UAS-dicer* control experiments. Genotypes are labelled below the graph.

**Fig. S15.**
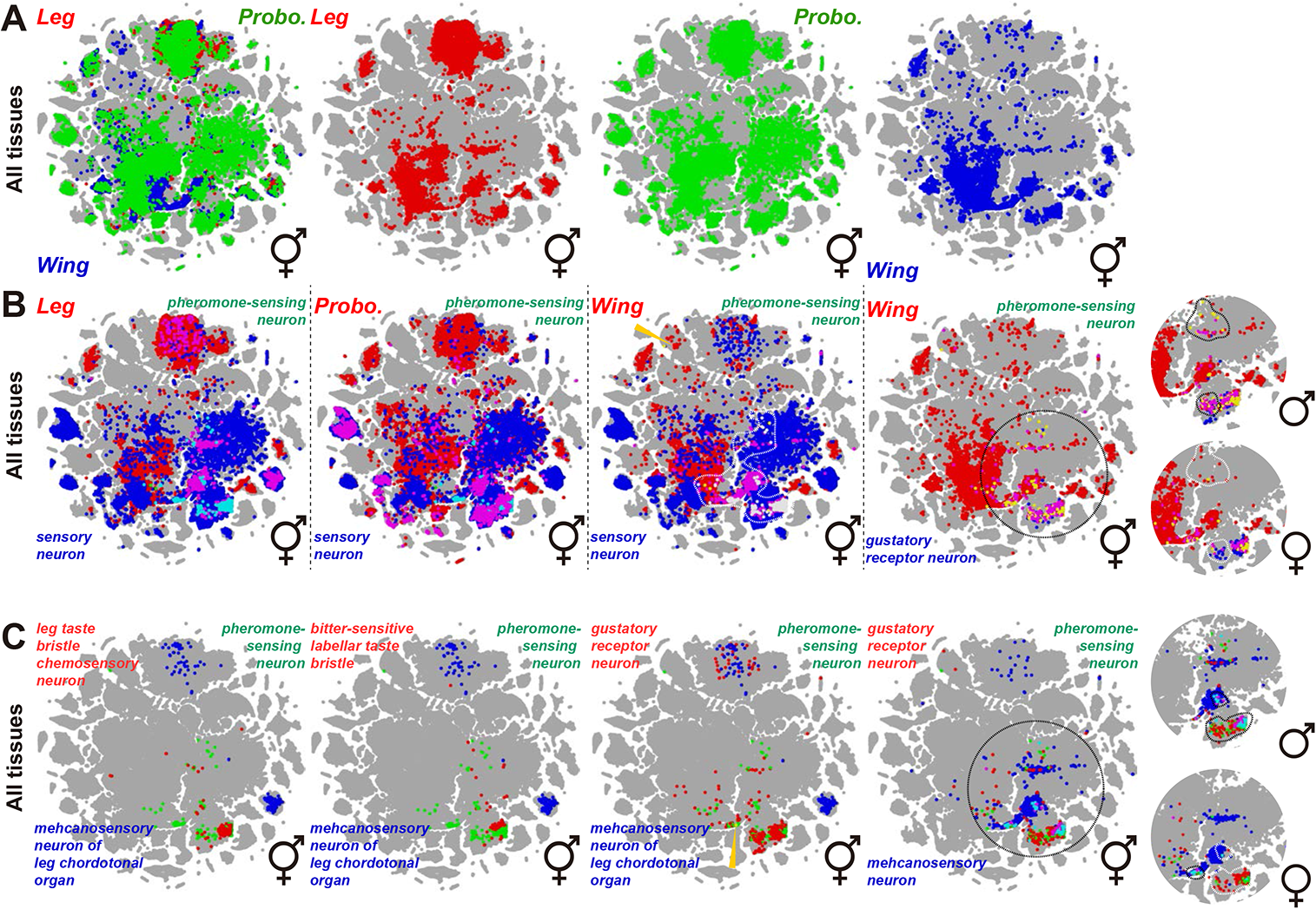
Whole tissue Fly SCope dataset. (A-C) Annotations and gene names are color-coded using red, green, and blue words. When cells overlap, the color of the dots is either yellow, cyan, or magenta. Colored arrows indicate cells that have been double- or triple-labeled by annotations and genes.

**Fig. S16.**
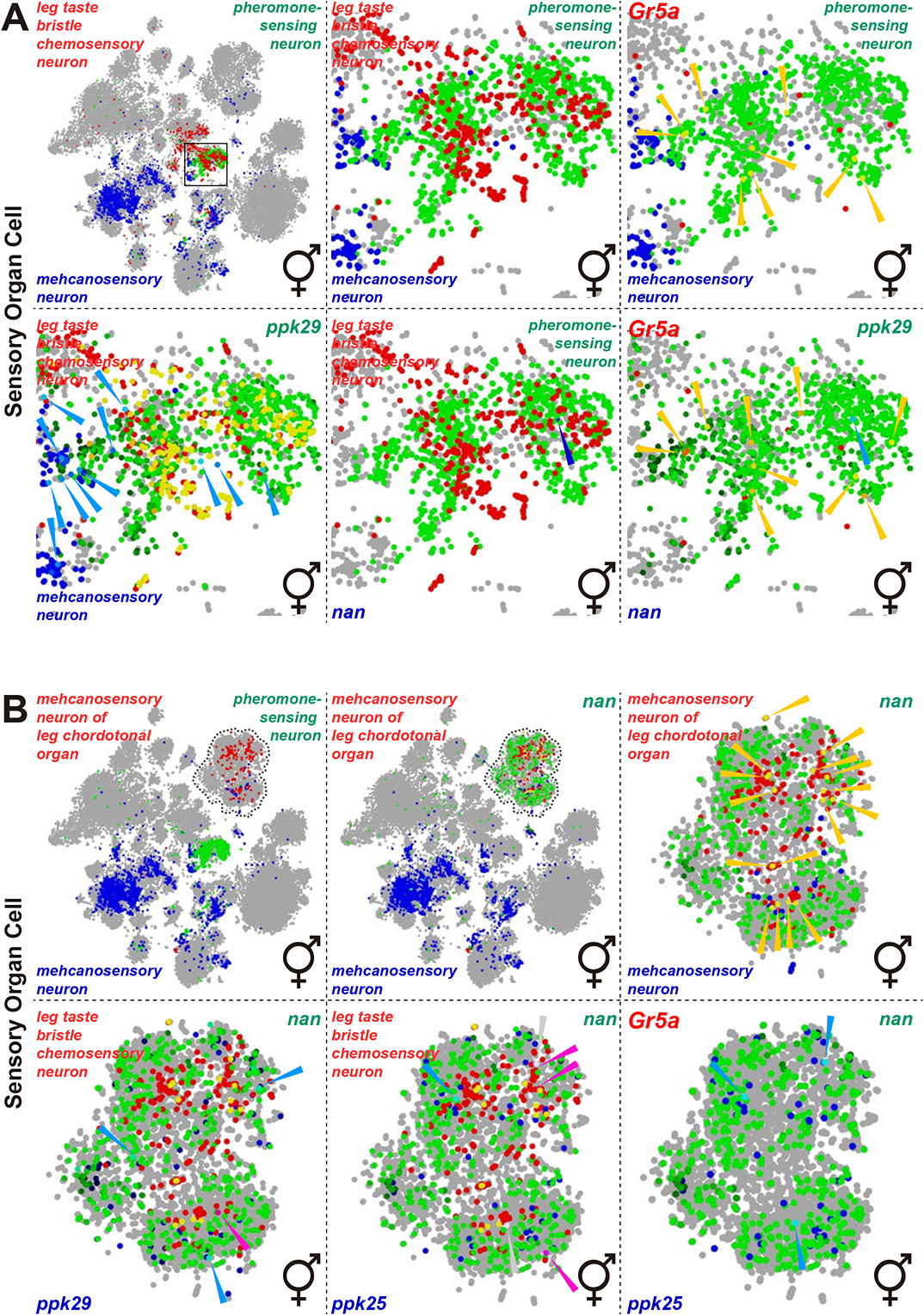
Sensory organ cell-specific Fly SCope dataset. (A-C) Annotations and gene names are color-coded using red, green, and blue words. When cells overlap, the color of the dots is either yellow, cyan, or magenta. Colored arrows indicate cells that have been double- or triple-labeled by annotations and genes.

**Fig. S17.**
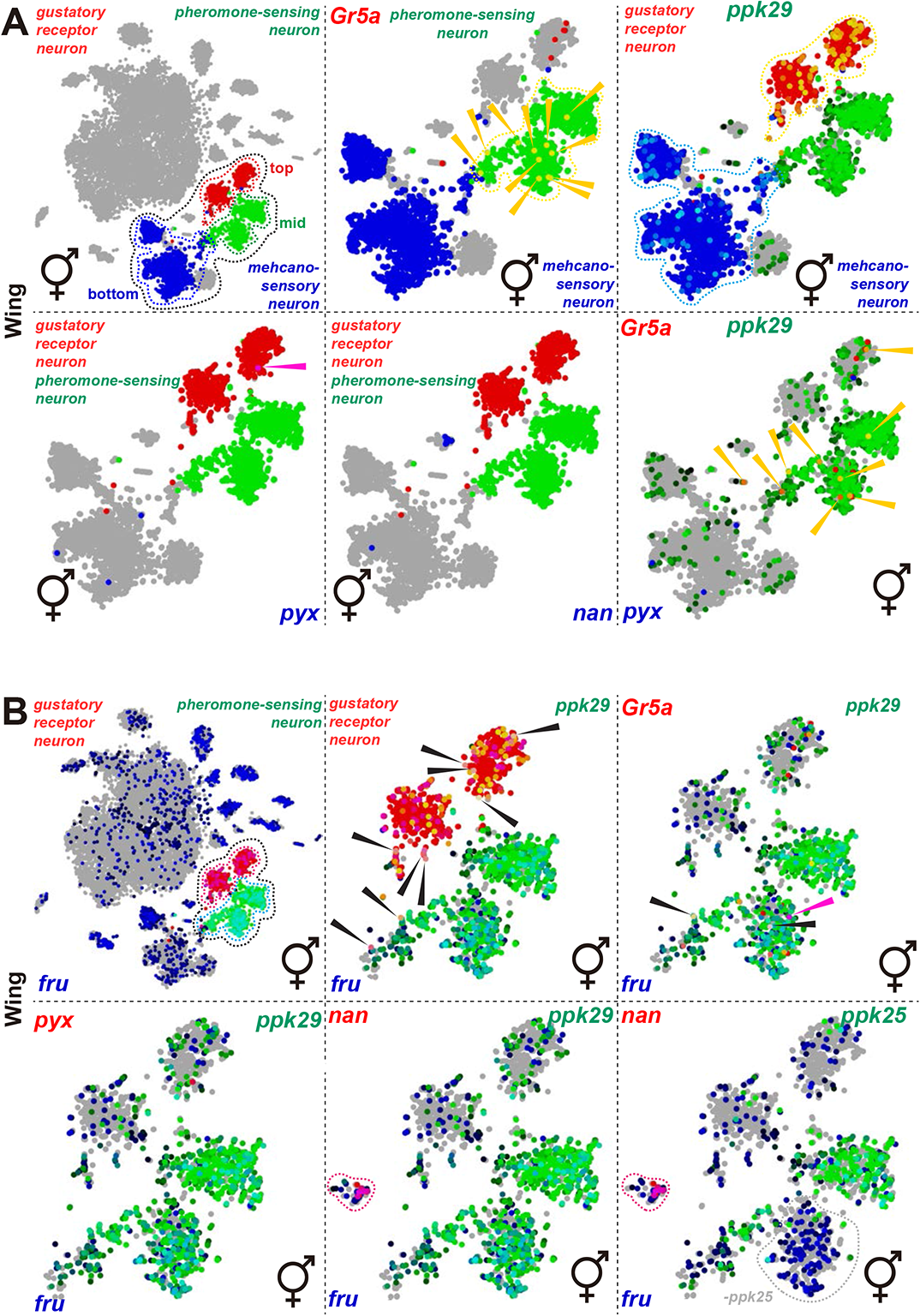
Wing-specific Fly SCope dataset. (A-C) Annotations and gene names are color-coded using red, green, and blue words. When cells overlap, the color of the dots is either yellow, cyan, or magenta. Colored arrows indicate cells that have been double- or triple-labeled by annotations and genes.

### Box. S1.

Costs and benefits of mate guarding. Here, we developed a mathematical model to identify the conditions under which experienced males gain fitness benefits by reducing mating duration compared to naïve males. The aim of this model is to generate testable evolutionary predictions to explain the observed SMD in experienced male *D. melanogaster*. In this model, we assume that i) males have a control of mate guarding duration and adaptively modify this duration to maximize their fitness benefits, and ii) the (reproductive) status of females are identical. We also assume that the costs of mate guarding increase linearly at the same rate in pre- and post-ejaculation period. Hence, without loss of generality, we consider that both the benefits and costs accumulate from the moment of sperm ejaculation for simplicity.

First, let us assume that the costs of mate guarding *C* accumulate linearly with time *t*. Then the costs are simply:

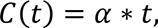

where *a* governs the rate of increase in the costs (*α* > 0). Let us also assume that the reproductive benefits of mate guarding increase asymptotically with a maximum threshold *β* (a maximum number of eggs a male can fertilize in a single mating, *β* > 0). Then the benefits of mate guarding *B* can be expressed as:

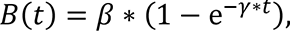

where *γ* determines how rapidly the benefits reach the maximum threshold (*γ* > 0). The benefits reach the maximum threshold faster when *γ* gets larger and *vice versa*. The net benefit of a male fly is then

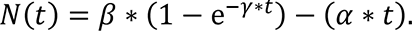

Because the second derivate of *N(t)* is always negative, *N(t)* has one optimal value of *t\** that maximizes it. Optimal mating duration *t\** is then

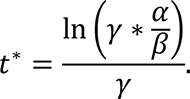

As long as *γ* > *α*/*β*, there exists one positive value of *t\**. From this equation, we can readily deduce that the optimal mating duration *t\** gets shorter when *α* gets larger (Fig. 12D), and/or *β* gets smaller (Fig. 12A). The relationship between *γ* and *t\** is not straight-forward, but dependent on the other parameters. If 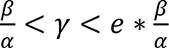, then *t\** get shorter as *γ* gets larger (Fig. 12B), but if 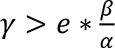, then *t\** gets shorter as *γ* gets smaller (Fig. 12C).

